# Defining Essential Enhancer for Pluripotent stem cells using Features Oriented CRISPR-Cas9 Screen

**DOI:** 10.1101/839316

**Authors:** Hao Fei Wang, Tushar Warrier, Chadi EL Farran, Zheng Zihao, Qiao Rui Xing, Melissa J Fullwood, Li-Feng Zhang, Hu Li, Jian Xu, Tit-Meng Lim, Yuin-Han Loh

## Abstract

Cis Regulatory Elements (CREs) regulate the expression of the genes in their genomic neighborhoods and influence cellular processes such as cell-fate maintenance and differentiation. To date, there remain major gaps in the functional characterization of CREs and the identification of its target genes in the cellular native environment. In this study, we performed a Features Oriented CRISPR Utilized Systematic (FOCUS) screen of OCT4-bound CREs using CRISPR/Cas9 to identify functional enhancers important for pluripotency maintenance in mouse ES cells. From the initial 235 candidates tested, 16 CREs were identified to be essential stem cell enhancers. Using RNA-seq and genomic 4C-seq, we further uncovered a complex network of candidate CREs and their downstream target genes, which supports the growth and self-renewal of mESCs. Notably, an essential enhancer, CRE111, and its target, *Lrrc31*, form the important switch to modulate the LIF-JAK1-STAT3 signaling pathway.

## INTRODUCTION

Cis-regulatory elements(CREs) are regions of non-coding DNA that regulate the expression of their target genes. 11% of the mouse genome was predicted to be non-redundant cis-regulatory elements (Shen et al., 2012). CREs were found to play roles in governing cell identity by regulating cell-type specific transcriptomic profiles (Buecker et al., 2014; Shen et al., 2012). Over the years, detailed identification and characterization of cell-type specific CREs were made possible through collaborative efforts such ENCODE and the Roadmap epigenomic project. Its hallmark includes the ability to regulate gene expression independent of their orientation and distance away from the target genes. Furthermore, a single CRE may regulate the expression of several genes at any one time or target different downstream genes in different cell types (Shlyueva et al., 2014). Notably, the major gaps in our knowledge of the cis-regulatory elements are the functional characterizations and the identifications of their target genes in the cell type of interest.

Many genetic approaches, such as reporter assays and STARR-seq (Arnold et al., 2013), were developed to address this. However, these methods relied heavily on the functional readout of the enhancer fragment outside of their native genomic architecture, which led to inaccurate representations of their endogenous activity. To fully address the contribution of cis-regulatory elements to biological systems within their genomic environment, it is imperative to disrupt their activities *in situ* within their genomic environment.

Pluripotency is the ability of stem cells to differentiate into all other cell types that constitute the entire organism. In the past few decades, many studies have defined the essential genes involved in maintaining pluripotency. Among them, *Oct4* was identified as a master transcription factor for the regulation of pluripotency and self-renewal in embryonic stem cells. The genomic binding profile of OCT4 protein in both the mESCs and hESCs has been elucidated (Chen et al., 2008; Loh et al., 2006; Boyer et al., 2005). Interestingly 30% of OCT4 bound sites were mapped to the distal regions (10-100kb) of the nearest gene, or to the gene deserts (>100kb to the nearest gene) (Loh et al., 2006). Recently several studies have been published on the characterization of CREs centered around the genomic regions of the *Oct4* gene (Diao et al., 2016; Diao et al., 2017).

Here, we describe a Features Oriented CRISPR Utilized Systematic (FOCUS) screen for essential OCT4-bound sites in mouse ES cells. From the screen, we identified seventeen high-confidence cis-regulatory elements which are critical for the maintenance of ES cells. Using a system approach, integrating genomic and functional analyses, such as ChIP-seq, ATAC-Seq, RNA-Seq, HiC-Seq, 4C-Seq, genetic knock-down, proteomics and rescue experiments, we further defined the target genes for these CREs and uncovered several novel regulators including the LRRC31-JAK-STAT3 axis, which plays a critical role in governing the proper signal transduction in ES cells.

## RESULTS

### FOCUS identified essential cis-regulatory elements for pluripotency maintenance

In order to functionally dissect the Oct4 bound cis-regulatory elements (CREs) that are essential for pluripotency maintenance in mouse ES cells, a Features Oriented CRISPR Utilized Systematic (FOCUS) screen of OCT4-bound CREs was conducted. To build the FOCUS library (Figure 1A, **Figure S1A**), we first collated mESC ATAC-seq data to ascertain the accessible chromatin regions(Li et al., 2017). Out of 27,513 sites identified, 6,392 demonstrated high confidence OCT4-binding based on published ChIP-Seq datasets (**Table S1**) (Chen et al, 2008). Using FIMO (Grant et al., 2011), we next determined intergenic OCT4-bound CREs, which contain the *Oct4* motif. Finally, we shorted listed putative targets for the primary screen after taking into account the presence of PAM motif (NGG) in or near the CREs.

**Figure 1.**
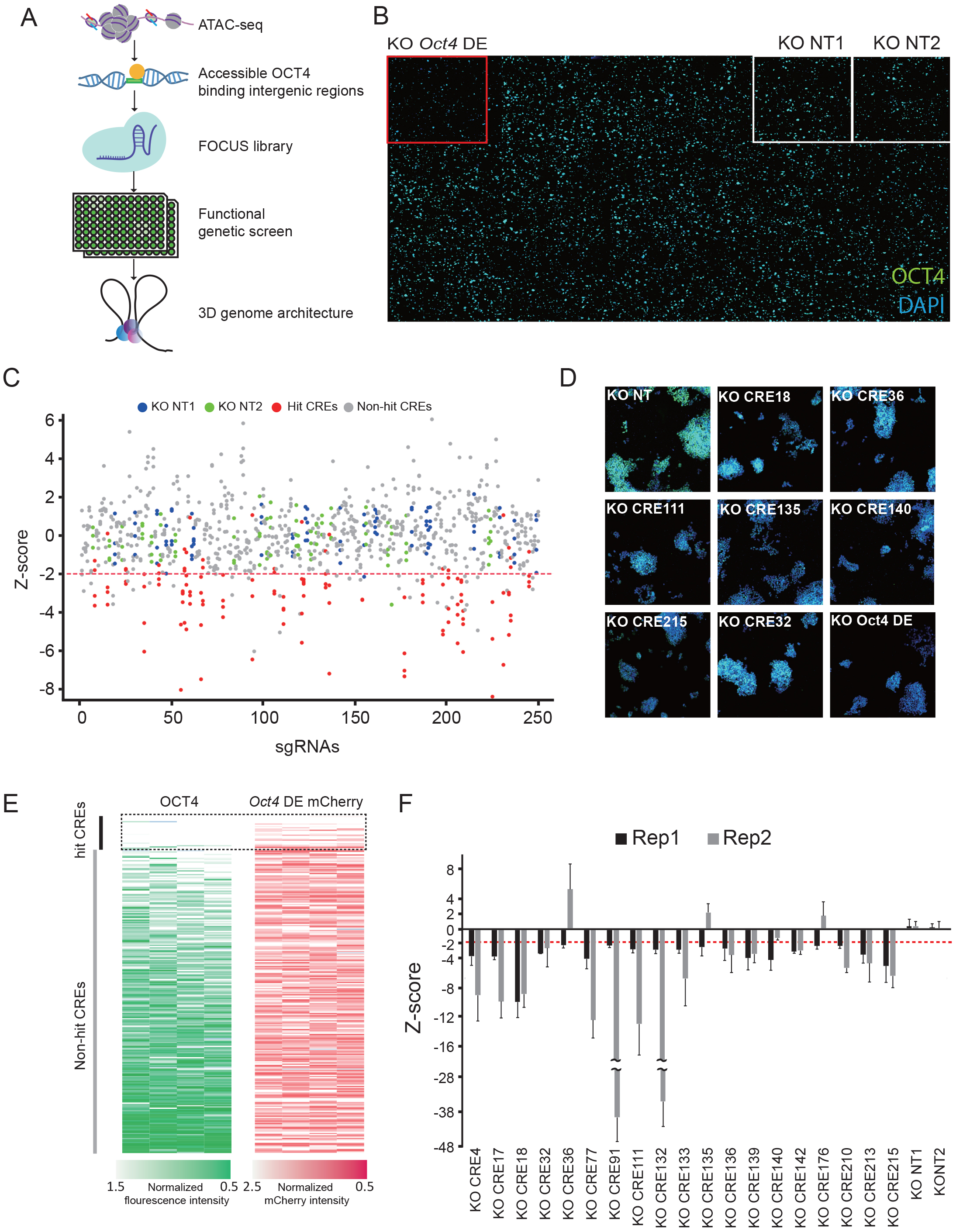
FOCUS screen identified cis-regulatory elements essential for pluripotency maintenance. (A) Schematic of FOCUS screen. sgRNAs targeting accessible OCT4 binding cis-regulatory elements in mESCs were designed and transfected into mESCs for the FOCUS screen. (B) An image of the FOCUS screen. sgRNA targeting the Oct4 Distal Enhancer was used as positive control (red square). KO NT1 and KO NT2 were used as negative controls (white squares). OCT4 immunofluorescence signal and nuclei staining are shown in green and blue respectively. (C) Distribution plot of the FOCUS screen. 19 out of 225 cis-regulatory elements with Z scores of <-2 (red dotted line) were identified as candidates potentially important for pluripotency maintenance (red dots). Non-targeting controls are labelled as blue and green dots. Remaining cis-regulatory elements are marked in grey. Z-score of 4 replicates are shown in the same line for each cis-regulatory element. (D) Representative images of OCT4 (green) and DNA (blue) for selected hits identified from the FOCUS screen. (E) Primary screen of 225 cis-regulatory elements with two independent assays, OCT4 (green) and Oct4-DE *mcherry* reporter (red). Each row represents the knockout of a single cis-regulatory element. Each column represents individual replicates. (F) Secondary screen of 19 candidate CRE hits. Rep 1 and 2 are two sgRNAs targeting different locations of the same cis-regulatory element. Z-score was calculated with a reference to KO NT2. The bar chart shows mean ± SD of 3 biological replicates.

sgRNAs were designed for 235 CREs based on their targetability by the CRISPR/Cas9 genome editing system, as assessed by the CRISPR sgRNA designer tool (**Table S2**) (Doench et al, 2016). Two non-targeting sgRNAs (NT1 and NT2) were used as negative controls in the FOCUS screen, whereas sgRNA against the Oct4 distal enhancer (DE) was used as a positive control (Figure 1B). Two independent read-outs, namely the Oct4 immunofluorescence signal and the *Oct4*-DE mCherry reporter signal, were used. We validated the specificity of the *Oct4*-DE mCherry reporter by comparing its activities in E14 mESCs, differentiated E14 and MEF cells (**Figure S1B, S1C and S1D**). To eliminate the bias introduced by the number of cells in our screens, we first assessed the correlation between cell number and OCT4 immunofluorescence signal in mESCs (**Figure S1E**). To this end, the correlation was used to normalize OCT4 immunofluorescence signal derived from both the primary and secondary screens. For each of the 235 cis-regulatory elements, z-score was calculated from 4 biological replicates (Figure 1C). Based on a targeted error rate of 0.05, a cut-off threshold of absolute value >2 SD from the mean of the non-targeting controls was used to determine candidate CREs which resulted in the reduction of OCT4 signal when they were knocked out (KO). Expectedly, the control sgRNA targeting the *Oct4* DE was the top hit from the screen with an average z-score of -4 (Figure 1C, **Table S3**). The observation that the average z-scores of both non-targeting sgRNAs fell within -2 to +2 further increased the confidence of our FOCUS screen (Figure 1C). To exclude the error introduced during the imaging analysis, random images of hits were extracted to confirm the reduction of OCT4 immunofluorescence signal (Figure 1D). To further test the robustness of our screen, the result was independently normalized to the two non-targeting sgRNAs, and the correlation between these two sets of analyses was calculated. We observed a Pearson correlation coefficient of 0.9 (**Figure S1F**). We next examined the *Oct4* DE mCherry signal for the KO CREs. We detected good correlation between OCT4 immunofluorescence and *Oct4* DE mCherry signal for both the hit and non-hit CREs (Figure 1E). To confirm their knock-out, the targeting efficiency of 10 randomly picked hit CRE sgRNAs were examined using the SURVEYOR assay (**Figure S1G**). For three CREs, we evaluated the percentage of mutated *Oct4* motif in the sgRNA-transfected mESCs using DNA sequencing. Around 38% to 47% of the sgRNA-transfected mESCs showed DNA deletion at the targeted *Oct4* motif (Figure S1H). Further, to exclude off-target effects of the sgRNAs, a secondary screen was performed using independent sgRNAs targeting the 19 hit CREs identified from the primary screen. Of note, 16 candidate CREs showed consistent phenotypes from the two batches of sgRNA KO (Figure 1F).

### Knockout of candidate CRE hits affects both the maintenance and establishment of pluripotent stem cells

A detailed evaluation of the deleterious effects of knocking out the candidate CREs was then undertaken. First, to examine the consequence of our CREs on self-renewal of mESCs, we assayed colony formation in cells in which candidate CREs were knocked out. For seven CREs, a marked decrease in the ability of the cells to form ES-like colonies was observed, based on a final colony count after alkaline phosphatase (AP) staining (Figure 2A). We detected similar phenotypic outcome when CREs were knocked out in ES-D3 cells using a different set of sgRNAs (**Figure S2A**). Next, the expression of a set of pluripotency and differentiation genes was determined by qRT-PCR upon knockout of candidate CREs in ES-E14 cells. As expected, the *Oct4* expression level decreased significantly when candidate CREs were knocked out (Figure 2B). Interestingly, we did not observe comparable downregulation of *Nanog* and *Sox2* across the different CREs KO. For most CREs, a universal increase in the expression of differentiation markers was detected with no tendency towards any particular lineage. Notably, we observed a specific upregulation of *Gata6* for KO CRE 4, suggesting that individual CREs may regulate pluripotency through varied pathways. We then induced differentiation towards the ectoderm lineage in ES-E14 cells by treatment with retinoic acid, followed by the measurement of the expression of several marker genes (*Pax6*, *Gbx2*, *Foxj3*, *Mcm7* and *Sox1*) (Zhang et al., 2015). Notably, knockout of the candidate CREs led to an increase in the expression of ectoderm markers, suggesting that the loss of function of our candidate CREs could destabilize the pluripotency state of ES cells and prime the mES cells for directed differentiation (Figure 2C, **S2B**).

**Figure 2.**
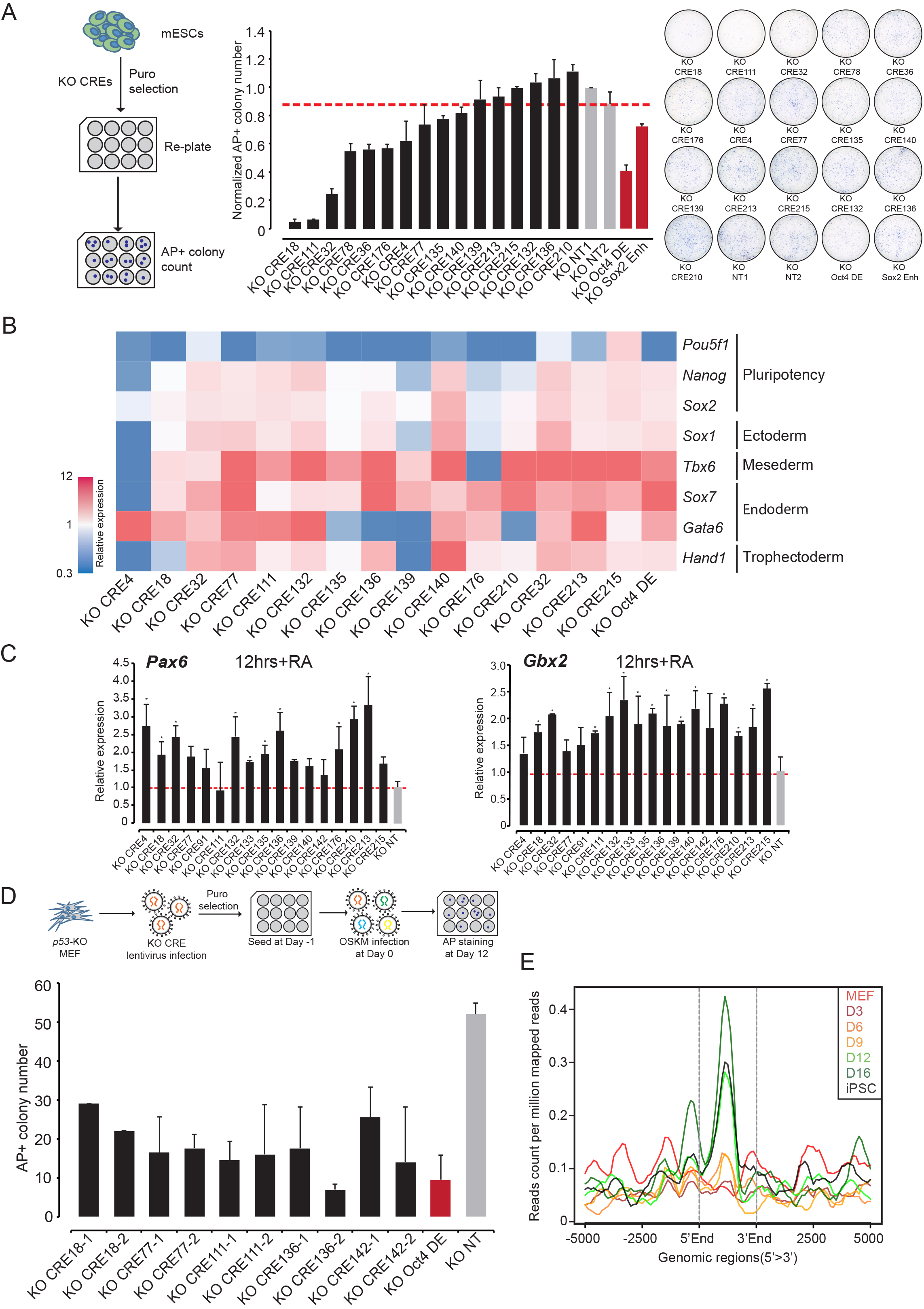
Knockout of candidate CRE hits affects both the maintenance and establishment of pluripotent stem cells. (A) Colony formation assay showing that the knock-out of candidate CREs compromised self-renewal of mESCs. Left: Schematic of colony formation assay. Middle: Normalized AP+ colony number of CREs (black) knockout. KO NTs (grey) were used as negative controls, while KO Oct4 DE and KO Sox2 Enh (red) were used as positive controls. The red dotted line shows the level of normalized AP+ colony number of KO NT2. The bar chart shows mean ± SD of 3 biological replicates. Right: Representative images of each candidate CRE knockout. (B) qRT-PCR showing decreased *Oct4* expression and increased lineage-specific gene expression at Day 3 post knockout of candidate CREs. Each row represents the expression of a gene. Each column represents the knockout of an individual candidate CRE. Data shown was normalized to E14 transfected with KO NT sgRNA. (C) qRT-PCR showing increased expression of differentiation genes when candidate hit CREs were knocked out during RA-induced differentiation. Cells transfected with KO NT sgRNA was used as control (grey) and all data was normalized to it. The bar chart shows mean ± SD of 3 replicates. Student’s t-test was used for statistical analysis. * represents p-value <0.05. (D) Knockout of candidate CREs affected the establishment of pluripotency during somatic cell reprogramming. Schematic of the experiment (top). Reprogramming efficiency was assayed at 10 d.p.i by AP staining (bottom). KO NT (grey) was used as a negative control, while KO Oct4 DE (red) was used as a positive control. The bar chart shows mean ± SD of 3 biological replicates. (E) Average enrichment plot of ATAC-seq signal of candidate CREs during somatic cell reprogramming. Different time points are shown in different colours.

Next, a somatic cell reprogramming assay was performed by infecting MEFs with the sgRNAs to knock out candidate CREs, before reprogramming them to induced pluripotency via the OSKM retroviral system (Takahashi and Yamanaka, 2006). Decreased reprogramming efficiency was seen for most candidate CREs as compared to the non-targeting control, when assessed by colony counting based on AP staining 10 days post-infection (d.p.i). (Figure 2D, **S2C**). In order to ascertain that the final phenotypic effects were independent of any proliferation defects, a cell proliferation assay was performed for these CREs KO MEFs. Only slight increase in cell proliferation rate was observed in most of the CREs KO MEFs compared to the KO NT. This suggests that the reduced reprogramming efficiency was indeed specific and not due to changes in cell proliferation (**Figure S2D**). We then examined the chromatin state of candidate CREs based on ATAC-seq datasets (Fang et al., 2018). For the hit CREs which affected the reprogramming process, we observed a gradual opening of its chromatin regions from Day 0 to Day 12 of reprogramming (Figure 2E, **S2E and S2F**). Together, the data suggests that the candidate CREs identified from the FOCUS screen play important roles in both the maintenance and establishment of pluripotent stem cells.

### Candidate CREs display enhancer activity in mESCs

We next investigated the profile of histone markers (H3K4me, H3K4me3, H3K27ac, H3K9ac, H3K9me3 and H3K27me3) on our hit CREs using published ENCODE datasets(ENCODE Project Consortium, 2012). A significant enrichment of active enhancer histone marks (H3K4me1 and H3K27ac) was detected in our candidate CREs (Figure 3A, 3B, **S3A, S3D and S3E**). Apart from histone modifications, a significant enrichment was also observed for pluripotency-associated transcription factors (OCT4, SOX2 and NANOG) and active enhancer-related transcription factors (MED1 and P300) (Figure 3C, **S3B, S3D and S3E**) (Chen et al., 2008). The enrichment of transcription factors’ motifs was further compared between the hit CREs and non-hit CREs. Notably, a specific enrichment of pluripotency-related transcription factor motifs (*Brn1*, *Zic3*, *Sf1* and *Arnt*) was detected in the hit CREs which reinforced the significances of multiple transcription factors co-binding on functional enhancers (Figure 3D, **S3C**) (Forristal et al., 2010; Fuellen and Struckmann, 2010; Gu et al., 2005; Lim et al., 2007; Kim et al., 2018). Of note, when *Zic3* or *Brn1* motif were mutated, the enhancer activity of CRE132 was further depleted, indicating the importance of these identified motifs (Figure 3E).

**Figure 3.**
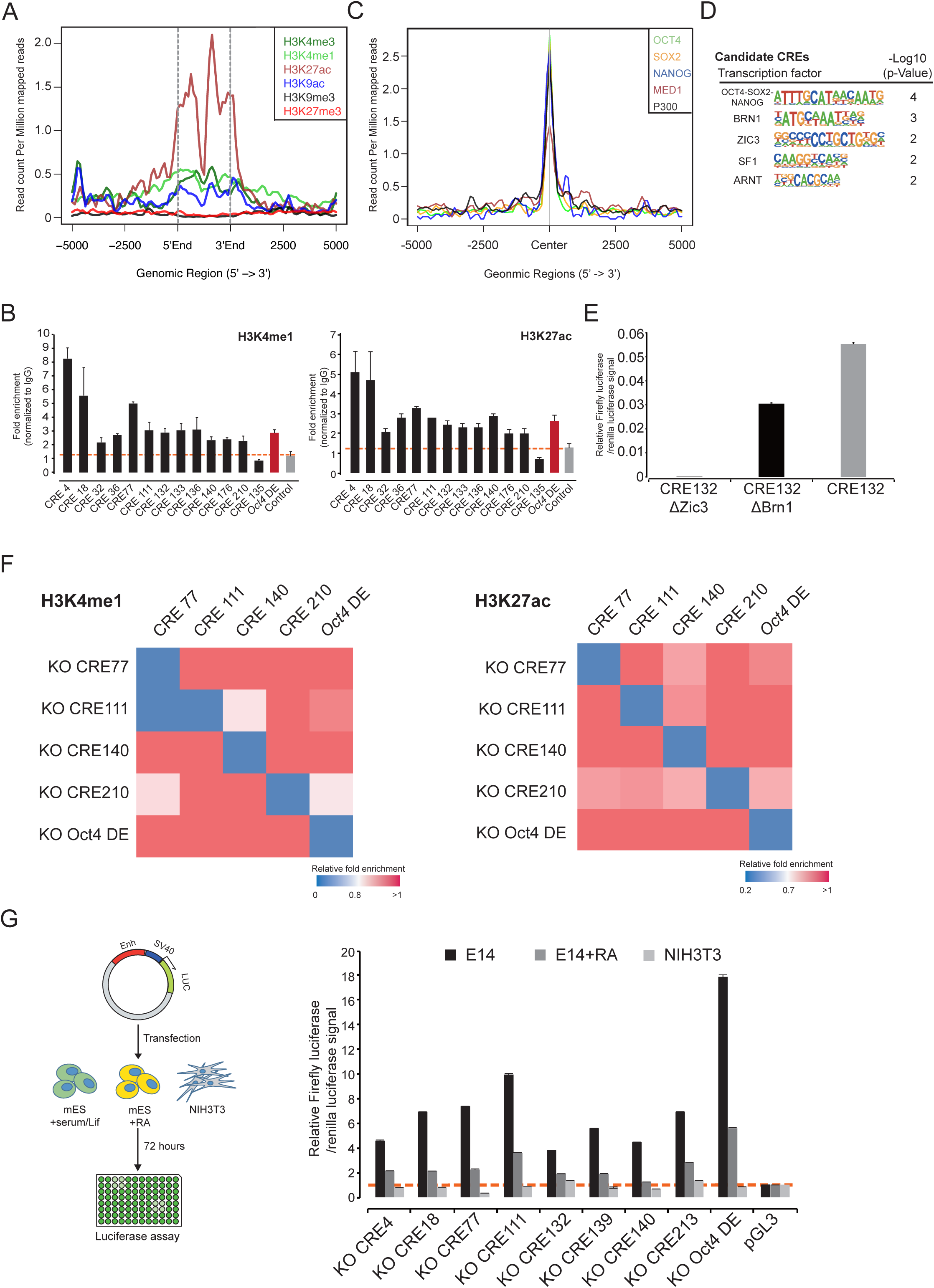
Candidate CREs display typical enhancer activity in mESCs. (A) Average enrichment plot of histone marks on hit CREs. Different histone marks are indicated in different colours. (B) Enrichment of H3K4me1 (left) and H3K27ac (right) on hit CREs (black) compared to a negative control region (grey) by ChIP-qPCR. Oct4 distal enhancer was used as positive control (dark red). The red dotted line shows the background enrichment of the negative control region. Data are representative of at least 3 independent experiments. The bar chart shows mean ± SD. (C) Average enrichment plot of pluripotency transcription factors on hit CREs. Different transcription factors are indicated in different colours. (D) Transcription factor motifs enriched at candidate CRE sites. (E) Luciferase activity of reporter plasmids containing CRE132 with mutated *Zic3* and *Brn1* motif. The bar chart shows mean ± SD of 3 independent experiments. (F) Knockout of *Oct4* motif decreased the enrichment of H3K4me1 (left) and H3K27ac (right) active histone marks on candidate CREs. Each row represents the relative enrichment of the histone modifications on five candidate CREs when one CRE was knocked out using sgRNA. The relative enrichment of histone marks is shown in a color scale ranging from blue (decreased binding) to red (no change as compared to control). (G) Luciferase activity of reporter plasmids containing a fragment of candidate CREs in E14 (black), RA treated E14 (dark grey) and 3T3 (light grey) (right). Schematic of the experiment was shown on the left. The bar chart shows mean ± SD of 3 independent experiments.

We next examined active enhancer histone marks, namely H3K4me and H3K27ac, in four candidate CREs disrupted by sgRNAs (Figure 3F). We observed that the enrichment of these histone marks, as well as OCT4 binding (**Figure S3H**), was significantly reduced. This suggests that the binding of OCT4 is essential for the maintenance of the active histone marks on enhancer regions. Notably the decrease was specific to the locus where individual CREs were knocked out (Figure 3F, **S3F and S3G**). Furthermore, we functionally assessed the enhancer activity of the CREs using a luciferase reporter assay. Consistent with the enrichment of active histone marks and transcription factors, our hit CREs displayed a significantly elevated luciferase signal in E14 mES cells as compared to the control NIH3T3 cells. Remarkably, the luciferase activities decreased when ES-E14 cells were induced to differentiate via the introduction of retinoic acid. This strongly suggests that the enhancer activities of our candidate CREs are specific to the pluripotent mESCs (Figure 3G, **S3I**). Taken together, our data indicates that a majority of the CRE hits have important functional roles in pluripotent cells as active enhancers.

### Knockout of candidate CREs elicited overlapping and distinct transcriptomic effects

To further evaluate the function of candidate CREs in ES cells, RNA-seq libraries were generated for each CRE knockout in ES-E14. All the RNA-seq libraries were of good quality (**Figure S4A, S4B**). As expected, *Oct4* expression was downregulated in all CREs KO RNA-seq libraries (**Figure S4C**). Gene ontology (GO) analysis was performed on both the up-regulated and down-regulated genes when candidate CREs were knocked out. Genes involved in cell differentiation, endodermal cell lineage and multicellular organism development were enriched in the up-regulated genes. We also detected enrichment of genes implicated in the activation of MAPK activity and positive regulation of ERK1 and ERK2 cascade, supporting the notion that the knockout of hit CREs could prime ES cells for differentiation process (Figure 4A) (Lanner and Rossant, 2010). Amongst the down-regulated genes, a significant enrichment for genes involved in stem cell population maintenance was observed (Figure 4A). Apart from that, we also detected an enrichment for the WNT signalling pathway which has been reported to be involved in the maintenance of pluripotency in mESCs (Miyabayashi et al., 2007; Sokol et al., 2011). Using CTEN analysis, we uncovered the enrichment of different lineage-associated genes when candidate CREs were knocked out (Figure 4B).

**Figure 4.**
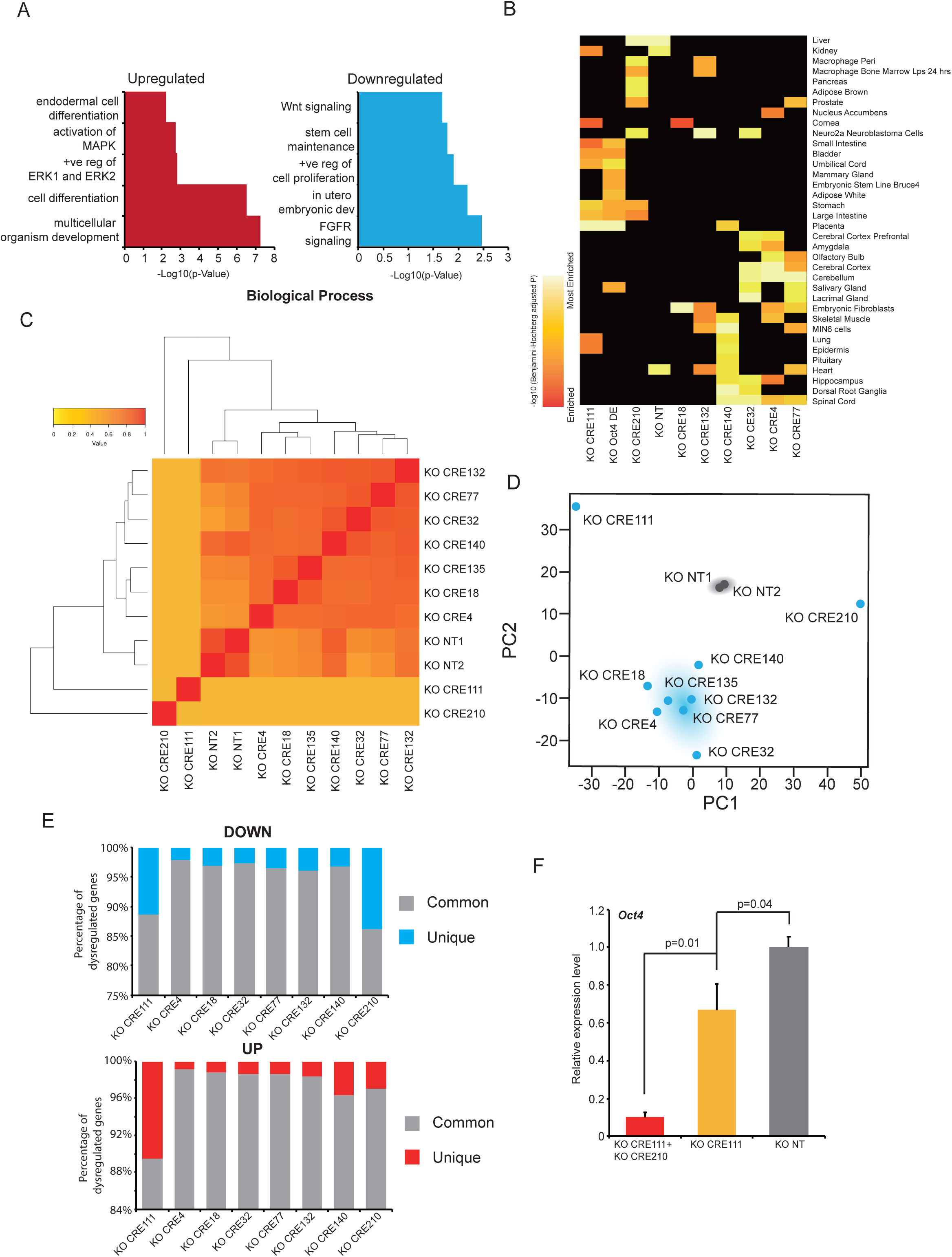
Knockout of candidate CREs elicited overlapping and distinct transcriptomic effects. A) Gene Ontology analysis of differentially expressed genes in the CRE KO libraries. X-axis represents the –Log10(p-Value) and Y-axis represents GO term. B) Lineage enrichment analysis of the CREs KO RNA-seq libraries. Each row represents a single lineage. Each column represents individual CRE KO RNA-seq library. -Log10 (Benjamini-Hochberg adjusted P) is represented as colour from red (enriched) to yellow (most enriched) C) Pearson correlation analysis shows that CREs 111 and 210 clustered distinctly. The correlation score is represented as colour from yellow (low) to red (high) D) Principal component analysis (PCA) of the CRE KO RNA-seq libraries. CRE111 clustered distinctively from the other CREs. KO NTs are shown in grey, CREs are shown in blue. E) Stacked column plot showing the percentage of overlapped genes which were differentially expressed between CRE KO and *Oct4* DE RNA-seq libraries. F) Combinatorial KO of CRE111 and CRE210 further decreased the expression of *Oct4* compared to single KO. Cells transfected with KO NT sgRNA was used as control (grey) and all data was normalized to it. The bar chart shows mean ± SD of 3 replicates. Student’s t-test was used for statistical analysis.

Next, we performed Pearson correlation analysis and principal component analysis (PCA) on these RNA-seq libraries (Figure 4C, 4D, **S4E**). Surprisingly, CRE111 and CRE 210 appeared as two distinct clusters compared to the other CREs. KEGG pathway analysis of the genes downregulated in CRE111 and CRE210 showed an enrichment for signalling pathways involved in pluripotency maintenance (**Figure S4D**). To test the similarity between CRE111, CRE210 and *Oct4* DE, dysregulated genes were compared between each KO CRE RNA-seq library and the KO *Oct4* DE RNA-seq library (Figure 4E, **S4F**). Indeed, KO CRE210 and CRE111 seemed to elicit transcriptomic profiles more similar to the KO *Oct4* DE. Through combinatorial knockout of CRE111 and CRE210, we observed a further reduction in *Oct4* expression and increase in differentiation gene expression compared to single knockout, suggesting these two CREs could affect pluripotency through different mechanism (Figure 4F, S4H). When we compared CRE111 KO RNA-seq library with the *Oct4* KD microarray library, a significant overlap was also observed (**Figure S4G**) (Loh et al., 2006).

### Candidate CREs regulate novel and known pluripotency genes

Next, to investigate the 3D genomic regions interacting with the hit CREs, we performed 4C-seq on seven CREs which showed the strongest phenotype in our functional assay (Figure 2). Out of these obtained high quality 4C-seq libraries for 4 CREs, we managed to get 4C-seq libraries with good quality (Figure 5A, **S5A, S5B and Table S5**), we then integrated the 4C-seq targets with KO CRE RNA-seq libraries to validate the functional enhancer-promoter interactions (Figure 5B, **Figure S5E**). For two CREs, CRE111 and CRE132, their interaction to putative target genes based on 4C-seq were validated using a detailed 3C assay (**Figure S5D**). Of the four CREs we examined for 3D interactions, most of the interacting regions localized between 5kb and 50kb to the nearest transcription start site (TSS) (**Figure S5C**).

**Figure 5.**
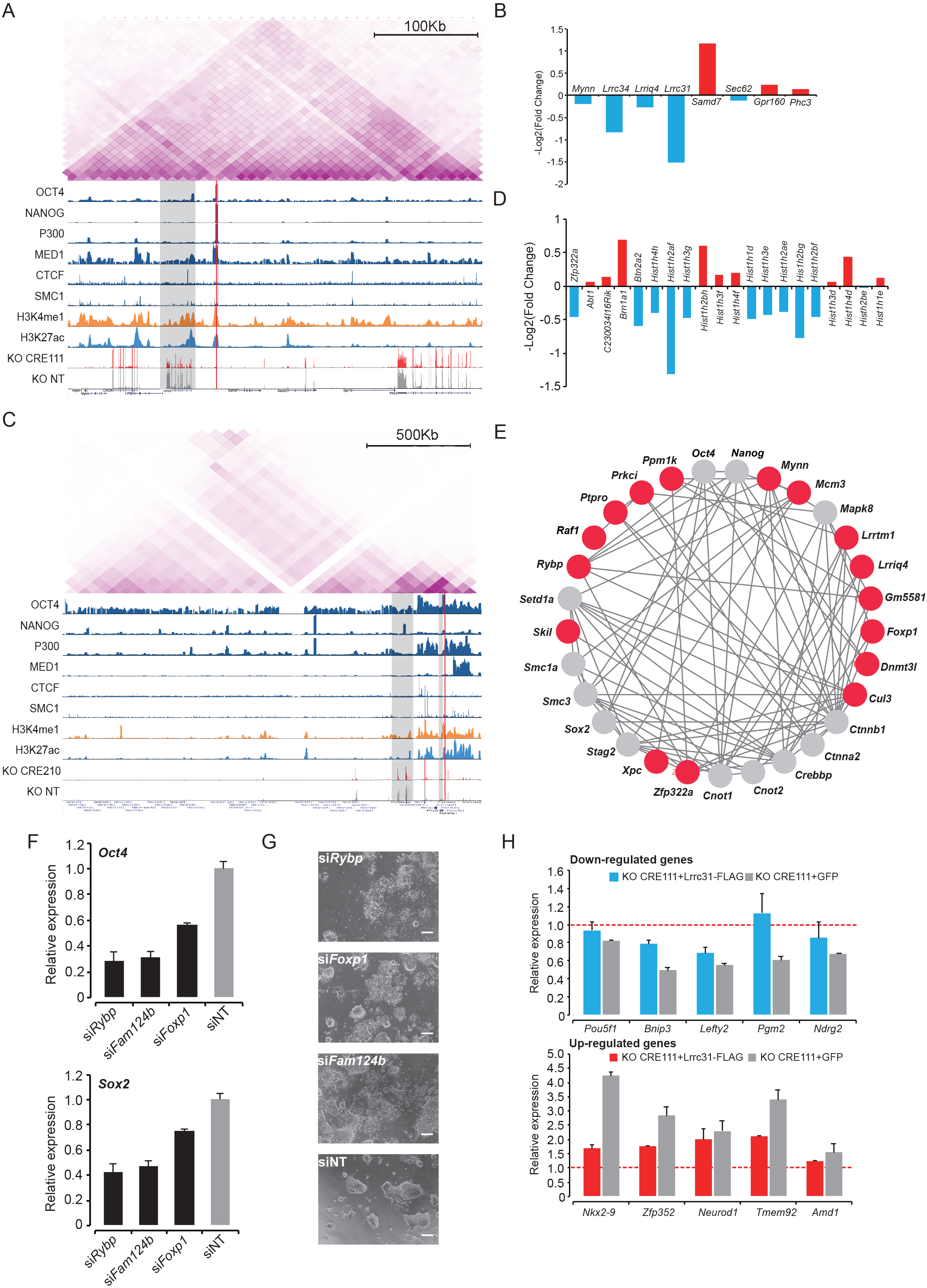
Candidate CREs regulate novel and known pluripotency genes. A) Integrative genomic view of Hi-C, ChIP-seq and RNA-seq data of CRE 111. Top panel: Hi-C interactions within the same TAD of CRE111. Middle panel: ChIP-seq UCSC browser of pluripotency and architectural proteins, as well as active enhancer marks. Bottom panel: RNA-seq UCSC browser of KO CRE111 and KO NT. The location of CRE111 is highlighted using red line. Putative target of CRE111 is indicated using grey shades. B) The bar chart showing the expression change of genes located within the same TAD as CRE111 when CRE111 was knocked out. The y-axis represents the -Log2(Fold change). The genes with decreased expression are shown as the blue bars. The genes with increased expression are shown as the red bars. C) Integrative genomic view of Hi-C, ChIP-seq and RNA-seq data of CRE 210. Top panel: Hi-C interactions within the same TAD of CRE210. Middle panel: ChIP-seq UCSC browser of pluripotency and architectural proteins, as well as active enhancer marks. Bottom panel: RNA-seq UCSC browser of KO CRE210 and KO NT. The location of CRE210 is highlighted using red line. Putative target of CRE210 is indicated using grey shades. D) The bar chart showing the expression change of genes located within the same TAD as CRE210when CRE210 was knocked out. The y-axis represents the -Log2(Fold change). The genes with decreased expression are shown as the blue bars. The genes with increased expression are shown as the red bars. E) Target genes of CREs form a tight network with known pluripotency regulators. Target genes are shown as red circles. Known pluripotency regulators are shown as grey circles. F) qRT-PCR showing that knockdown of CRE target genes (black) decreased the expression of *Oct4* (top panel) and *Sox2* (bottom panel) significantly as compared to siNT control (grey). The bar chart shows mean ± SD of 3 replicates. G) Brightfield images showing morphology changes in E14 cells induced by knockdown of CRE target genes. H) The interaction of CRE 111 and *Lrrc31* in E14 cells was confirmed by the 3C assay. X-axis shows 3C fragments corresponding to actual genomic locations. Y-axis shows relative interaction frequency between E14 and the BAC control. The corresponding location of each fragment in the genome is indicated using red dotted line. The bar chart shows mean ± SD of 3 replicates. I) qRT-PCR showing gene expression in the rescue of the expression of upregulated (red) or downregulated (blue) genes in CRE 111 KO, by the overexpression of LRRC31 in E14 cells. Red dotted line represents the relative expression of genes in WT E14 mESCs. The bar chart shows mean ± SD of 3 replicates.

For the remaining CREs, we performed an integrative analysis based on the published Hi-C dataset (Bonev et al., 2017) and our KO CRE RNA-seq libraries to predict the interacting gene targets (Figure 5C, **S5F and S5G**). Consistent with previous reports that topologically associating domains (TADs) are the boundaries for enhancer-promoter interactions, most of the interactions of candidate CREs happened within a TAD (Figure 5A, 5C **and S5F**) (Dixon et al., 2012). Interestingly, for CRE210, we identified *Zfp322a* (Ma et al., 2014), a known pluripotent gene, as its putative target. This further support the notion that our candidate hit CREs are critical to pluripotency maintenance (Figure 5C, 5D **and S5E**). Interestingly, potential interacting target genes of our candidate CREs formed a tight protein-protein network with known pluripotency regulators (Figure 5E). Furthermore, when we knocked down the target genes using siRNA, we observed significant decrease in the expression of pluripotent markers such as *Oct4* and *Sox2*, and a differentiation morphology as compared to siNT control (Figure 5E, 5F).

### *Lrrc31* regulates the maintenance of pluripotency in mES cells through the JAK-STAT3 signaling pathway

We next focused on one of the hit CREs, CRE 111, to further elucidate its detailed underlying mechanism in pluripotency maintenance. Firstly, to understand the functional significance of the interaction between CRE111 and its putative target gene, *Lrrc31*, we examined its over expression upon CRE111 knockout. Indeed, we observed partial rescue of the CRE111 KO dysregulated genes (Figure 5H). Hence, CRE111 may function through its activation of *Lrrc31* to govern stem cell pluripotency.

To address the role by which *Lrrc31* effects pluripotency in mESCs, we first knocked down *Lrrc31* using either siRNA or shRNA. Both assays lead to a differentiation morphology when *Lrrc31* was depleted (Figure 6A). As seen in the CRE 111 knock-out mESCs, *Oct4* expression was similarly decreased in *Lrrc31* depleted cells (**Figure S6A**), while the expression of differentiation genes was markedly upregulated (Figure 6B). Apart from knocking down *Lrrc31*, we also generated *Lrrc31* KO clones using CRISPR (**Figure S6D**). Compared to NT clone, *Lrrc31* KO leads to a significant reduction of pluripotent genes’ expression and self-renewal capacity of mESCs (Figure 6C, **S6E**). We next evaluated the role played by *Lrrc31* in mouse somatic cell reprogramming. Using siRNA, we depleted *Lrrc31* at different time points during reprogramming and observed a specific reduction of reprogramming efficiency when *Lrrc31* was knocked down at day 9 post viral infection (Figure 6D, **S6C**). Conversely, we observed drastic upregulation of *Lrrc31* in successfully reprogrammed cells compared to non-reprogrammed cells from our previously published RNA-seq datasets (**Figure S6F**) (Fang et al, 2018). Taken together, this data suggests that *Lrrc31* is an important effector for the acquisition of pluripotency at the late stage of reprogramming.

**Figure 6.**
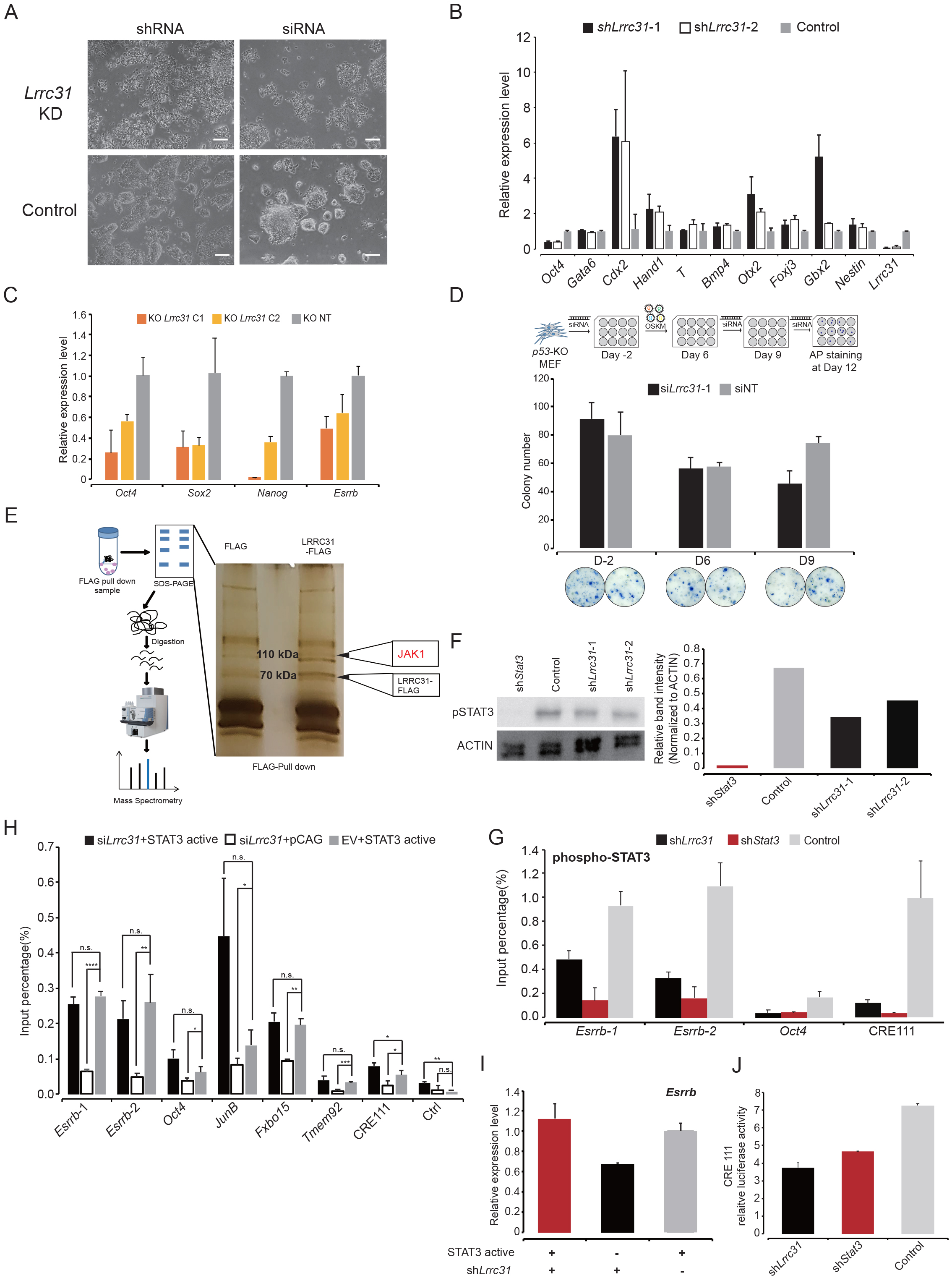
*Lrrc31* regulates the maintenance of pluripotency in mES cells through the JAK-STAT3 signaling pathway. A) Brightfield images showing the effects of knocking down *Lrrc31* using both siRNA and shRNA. Note the drastic change in morphology of the E14 cells. B) qRT-PCR showing that *Lrrc31* knock-down resulted in reduced expression of pluripotency genes and corresponding increase in expression of differentiation genes. Two constructs of *Lrrc31* shRNA are shown as dark and light blue bars, while the vector control is shown in grey. The bar chart shows mean ± SD of 3 independent experiments. C) qRT-PCR showing that knockout of Lrrc31using sgRNA reduced the expression of pluripotency genes. Two clones of *Lrrc31*-/- are shown as orange and yellow bars, while the *Lrrc31* +/+ is shown in grey. The bar chart shows mean ± SD of 3 independent experiments. D) Knockdown of *Lrrc31* at different time points of somatic cell reprogramming process. Top panel: Schematic of the *Lrrc31* knockdown during somatic cell reprogramming. Middle panel: Reprogramming efficiency (AP+ colony numbers) was assayed at 12 d.p.i. *Lrrc31* knockdown (black) at 9 d.p.i gave rise to reduced reprogramming efficiency as compared to the siNT control (grey). The bar chart shows mean ± SD of 3 biological replicates. Bottom panel: Representative images of each treatment group. E) Identification of LRRC31 interacting partners. Schematic of mass spectrometry experiment using LRRC31-FLAG pull down (left). Potential interacting partners of LRRC31 (right). Differentially enriched bands were highlighted with black arrows. The 75 KDa band represents the bait, LRRC31-FLAG protein. The 100KDa band represents potential interacting partners of LRRC31. The protein ID is listed in the box. High confidence hits are highlighted in red. F) Western blot shows that *Lrrc31* knockdown caused the corresponding decrease in STAT3 phosphorylation. Band intensity was quantified and normalized to ACTIN. G) ChIP qRT-PCR showing that the knockdown of *Lrrc31* decreased the binding of phospho-STAT3 on its targets in E14 cells. The shRNA of *Lrrc31* is shown as black bars. The empty vector of pSUPER was used as control (light grey). shRNA targeting *Stat3* was used as positive control (dark red). The bar chart shows mean ± SD of 3 independent experiments. H) Rescue of STAT3 binding on its target genomic regions by STAT3 dominant active mutant overexpression in *Lrrc31* knockdown ES-E14. The STAT3 dominant active mutant overexpression in *Lrrc31* knockdown ES-E14 is shown as black bars. The *Lrrc31* knockdown only is shown as white bars. The empty vector of pSUPER with STAT3 dominant active mutant overexpression was used as control (light grey). The bar chart shows mean ± SD of 3 replicates. I) Rescue of *Esrrb* expression by STAT3 dominant active mutant overexpression in *Lrrc31* knockdown ES-E14. The bar chart shows mean ± SD of 3 replicates. J) Relative luciferase activity of CRE111 reporter plasmids in sh*Lrrc31* transfected E14 (black), sh*Stat3* transfected E14 (red) and shEV transfected E14 (grey). The bar chart shows mean ± SD of 3 replicates.

To further elucidate its mechanistic role, we performed RNA-seq on *Lrrc31* knock-down mESCs. GO analysis on the upregulated genes showed enrichment for receptor binding function (**Figure S6I**). Consistent with qRT-PCR (Figure 6B), genes related to cell differentiation function were also enriched. A significant overlap between the *Lrrc31* knockdown and CRE111 knockout RNA-seq libraries was similarly observed, further supporting the notion that CRE111 controls pluripotency by regulating downstream *Lrrc31* (**Figure S6G**).

LRRC31 contains nine leucine-rich repeat domains, which was previously reported to function as a protein recognition motif (Kobe et al, 2001). We next overexpressed LRRC31-FLAG in mESCs and performed a mass spectrometry analysis to detect its binding partners (Figure 6E). A cytoplasmic localization was observed for LRRC31 in mESCs (**Figure S6B**). 18 proteins were identified as high confidence hits from two biologically independent samples performed at different proteomics facilities (**Table S6**). Among them, JAK1 was one of the top hits. Of interest, JAK1 has previously been reported to phosphorylate STAT3 in mESCs (Do et al., 2013; Onishi and Zandstra, 2015). Hence, we investigated the level of STAT3 phosphorylation upon *Lrrc31* knock down. A marked reduction in phosphorylated STAT3 was observed when *Lrrc31* was depleted (Figure 6F). To further assess the effect of *Lrrc31* on STAT3 function, STAT3 ChIP was performed on the *Lrrc31* knockdown cells. *Lrrc31* knockdown resulted in a significantly decreased binding of STAT3 on its target genes, including pluripotency factors such as *Oct4* and *Esrrb* (Figure 6H, **S6J, S6K**). Consequentially, decreased expression of STAT3 target genes in *Lrrc31* knockdown ES-E14 was observed (**Figure S6A**). Furthermore, a rescue of both the STAT3 binding and the expression of its target gene, *Esrrb*, was revealed when a dominant active STAT3 mutant was expressed in *Lrrc31* depleted ES-E14 (Figure 6H, 6I). Similar rescue was also observed for the *Lrrc31* KD-specific differentially expressed genes when dominant active STAT3 was overexpressed in *Lrrc31* KD mESC (**Figure S6L**). Interestingly, CRE111 was also bound by phospho-STAT3 and the binding was reduced in *Lrrc31* knockdown, suggesting a positive feedback regulation of *Lrrc31* expression (Figure 6G). This was further validated using luciferase assay of CRE111 when *Lrrc31* or *Stat3* was depleted in ES-E14 (Figure 6J).

Next, we hypothesized that the down-regulation of *Lrrc31* in F9 mouse embryonal carcinoma and hESCs, where LIF/JAK1/STAT3 signalling pathway is not essential, would not affect their respective cell state (Dahéron et al., 2004; Kawazoe et al., 2009). As expected, both knock down of *Lrrc31* and knock out of CRE111 in F9 cells did not affect the expression of pluripotency genes, such as *Oct4* (**Figure S6M and S6N**). Similarly, knock down of *Lrrc31* in hESCs elicited the same effects (**Figure S6O**). Taken together, our data suggests that *Lrrc31* functions as a novel regulator for pluripotency maintenance in mESCs by effecting the phosphorylation of STAT3 in mESCs.

### *Lrrc31* regulated the binding of STAT3 to the super-enhancers

To further elucidate the regulatory relations between *Lrrc31* and STAT3, we performed STAT3 ChIP-seq in mESCs when *Lrrc31* was knocked down by shRNAs. An enrichment of STAT3 motif was detected indicating the good quality of our STAT3 ChIP-seq libraries (Figure 7A). By overlapping the differential STAT3 binding peaks of two *Lrrc31* shRNA constructs, 4267 binding sites were chosen as the bona fide *Lrrc31*-regulated STAT3 binding sites (Figure 7B). GO analysis revealed that pluripotency-related signaling, such as TGF-beta and Wnt pathways, were regulated by *Lrrc31*(Figure 7C).

**Figure 7.**
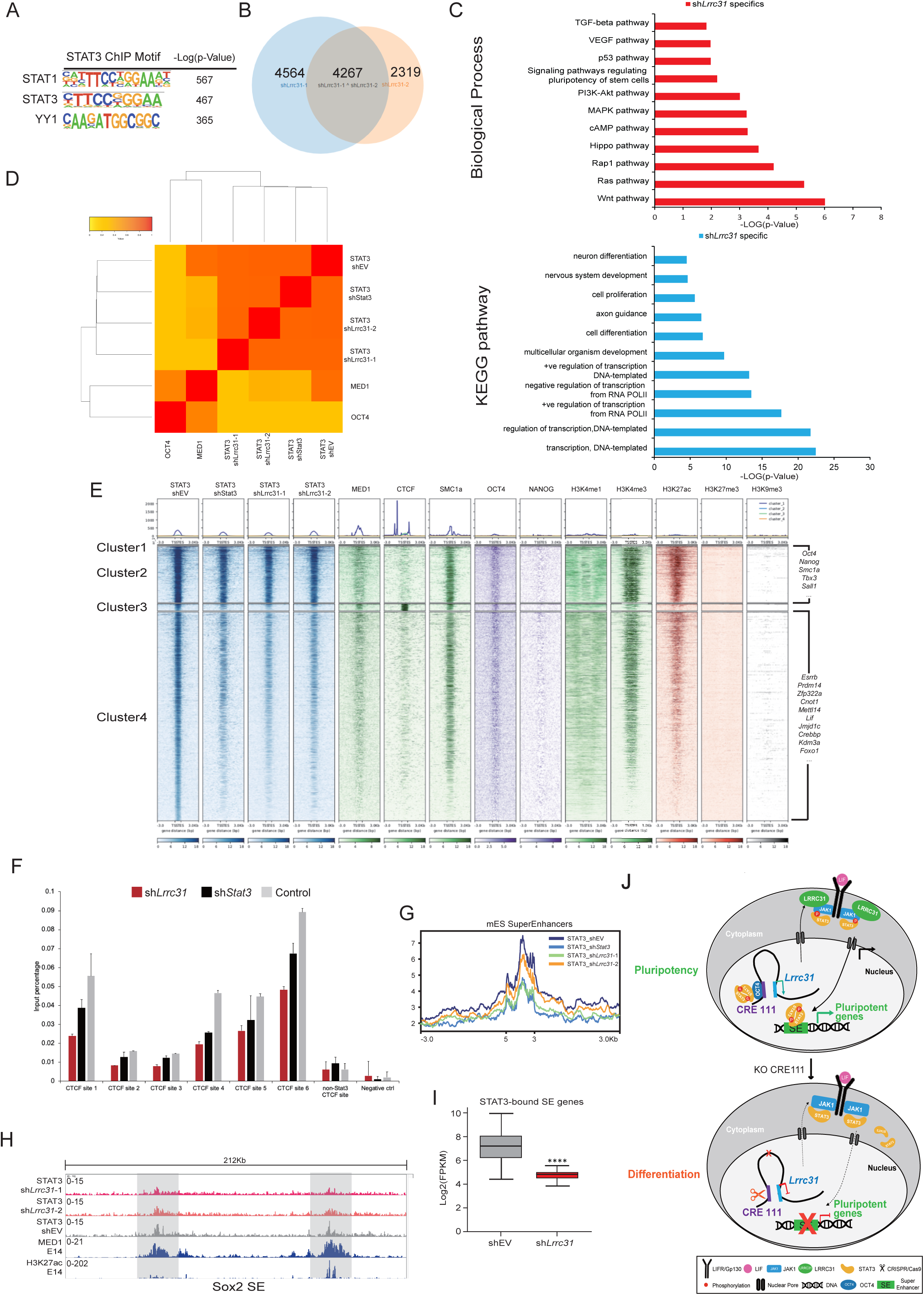
Lrrc31 regulated the binding of STAT3 to the super-enhancers. A) Transcription factor motifs enriched at STAT3 binding peaks. B) The Venn diagram showing the overlap of differentially STAT3 binding sites between two Lrrc31 shRNA constructs. C) Gene ontology analysis of differentially STAT3 binding sites when Lrrc31 was knocked down using shRNA. X-axis represents the –Log10(p-Value) and Y-axis represents GO term. D) Pearson correlation analysis shows the correlation of STAT3 binding peak between sh*Lrrc31* and sh*STAT3*. The correlation score is represented as colour from yellow (low) to red (high) E) Enrichment of several histone marks and transcription factors at the genomic regions of STAT3 binding sites. The heatmaps are clustered according to the enrichment profile of all the ChIP-seq libraries indicated in the figure. The top panel represent the average enrichment plot of the indicated ChiP-seq library. F) ChIP qRT-PCR showing that both *Lrrc31* knock-down and *Stat3* knock-down reduced the binding of CTCF on the cluster 3 STAT3 binding sites. The shRNA of *Lrrc31* is shown as red bars. The empty vector of pSUPER was used as control (light grey). shRNA targeting *Stat3* was shown as black bars. The bar chart shows mean ± SD of 3 independent experiments. G) Average enrichment plot showing the reduced binding of STAT3 on the super-enhancer when *Lrrc31* was knocked down. The y-axis represents average normalized number of fragments at the corresponding genomics regions indicated in he x-axis. H) UCSC screenshot of binding profile of STAT3 on Sox2 super-enhancer (indicated by the binding of MED1 and H3K27ac) when *Stat3* or *Lrrc31* was knocked down. I) Box plot showing the expression level of super-enhancer target genes when *Lrrc31* was knocked down by shRNA. Student’s t-test was used for statistical analysis. **** represents p-value <0.0001. K) In WT mESCs (top), OCT4 binds onto CRE 111 to maintain the expression of Lrrc31. LRRC31 interacts with JAK1 to phosphorylate STAT3 upon the binding of LIF on LIFR/Gp130. Phosphorylated STAT3 is then translocated into the nucleus and binds onto both pluripotent genes’ promoter, such as *Oct4* and *Esrrb,* and super-enhancers to activate their expression. Knocking out CRE111 using CRISPR (bottom) decreases the expression of Lrrc31. The loss of LRRC31 affects the phosphorylation of STAT3 through JAK1. Decreased level of phosphorylated STAT3 further diminishes the expression of downstream pluripotency genes.

Next, we clustered STAT3 ChIP-seq libraries with published transcription factors and histone modification ChIP-seq libraries (Figure 7D, 7E). Out of the four clusters, cluster 2 and cluster 4 showed quite a similar pattern with the decreased STAT3 binding and enrichment of active histone modifications, H3K4me3 and H3K27ac, SMC1 and OCT4 (Figure 7E). For both clusters, we detected the enrichment of pluripotent related genes. Interestingly, for cluster 3, we observed a strong enrichment for CTCF and the deprivation of active histone modifications, which is confirmed by the motif enrichment analysis (**Figure S7A**). To test whether the binding of STAT3 is essential for the binding of CTCF at the cluster 3 sites, we performed CTCF ChIP-qRT-PCR after the knock-down of *Lrrc31*, *Stat3* or control NT in mESCs. Interestingly, a reduction of the CTCF’s binding of cluster 3 sites was detected when either *Stat3* or *Lrrc31* was knocked down (Figure 7F).

Apart from CTCF, we also observed an enrichment of both MED1 and H3K27ac, markers for super-enhancer, on the cluster 2 and cluster 4 sites (Figure 7E). Using published super-enhancer dataset, we detected an enrichment of STAT3 on the super-enhancers of mESCs and the enrichment was reduced when *Lrrc31* was knocked down (Figure 7G, 7H, **S7B and S7D**). GO analysis on the STAT3-bound super-enhancers’ target genes revealed an enrichment of genes associated with stem cell population maintenance (**Figure S7C**). Moreover, these target genes’ expression was significantly reduced in *Lrrc31* knock-down mESCs (Figure 7I). Together, our data suggested that, *Lrrc31* actuates the binding of STAT3 on the super-enhancers of mESCs through its regulation of STAT3 phosphorylation (Figure 7J).

## DISCUSSION

Several recently-published studies sought to characterize and identify important CREs which may be involved in biological processes of interest (Diao et al., 2017; Fulco et al., 2016; Korkmaz et al., 2016). These screens utilized a pooled strategy involving the application of large number of CRE specific sgRNAs. Nonetheless, pooled screens have been reported to result in high false discovery rates, in part due to the introduction of biases at different stages of the screen. Whereas, our FOCUS screen pre-selected 235 CREs based on ATAC-seq and OCT4 ChIP-seq data in mESCs. Furthermore, each CRE was knocked out individually to assess its function in pluripotency. As a result, of the 19 CREs identified as putative hits from our primary screen, 84% could be validated in the secondary screen. This indicates the robustness and reliability of our method. To date, our study is the first systematic CRISPR screen to identify novel CREs, localized to the intergenic genomic regions, involved in the regulation of pluripotency.

Of note, we detected a large number of the candidate CREs which displayed enhancer activity in mESCs. This further highlights the key role that enhancer elements play in the maintenance of cell identity. Lending support to the observation was the enrichment of pluripotency transcription factors and proteins relating to enhancer function (eg. P300 and MED1) at our essential stem cell CREs. It is noteworthy that from the FOCUS screen, 5 candidate CREs did not exhibit characteristics typical of active enhancers, suggesting that they might regulate pluripotency in a different manner.

CRE 111, an essential stem cell enhancer was found to target and regulate *Lrrc31*. Using protein mass spectrometry and ChIP, we further observed that Lrrc31 functions through JAK1 to modulate the phosphorylation of STAT3. It is worth noting that the expression level of LRRC31 is maintained at basal level in mESCs and the overexpression of LRRC31 shows strong cytotoxicity in mESCs (data not shown). This suggests that LRRC31 expression in mESCs is fine-tuned within an optimal range for its normal cellular function.

As with all screens, our study has its drawbacks, including the limited number of targetable CREs. In our strategy, CRE KO is achieved by mutating the Oct4 motif inside the CRE using CRISPR Cas9. Based on published literature (Cong et al., 2013), Cas9 usually will generate indel from 1bp to 20bp through NHEJ. Thus, making the sgRNAs targeting within the 14bp Oct4 Motif is extremely important for the efficiency of CRE KO. But, due to the PAM domain restriction, a large number of CRE (364 CREs) with Oct4 motif do not have a targetable site within the motif. Thus, by using modulated Cas9 fusion proteins, such as dCas9-LSD1 or dCas9-KRAB, we can mitigate the restrictions and to increase the coverage of FOCUS screen. Similarly, use of Cas variants with alternative PAM domain, would enable the targeting of more CREs with less sequence restriction (Kearns et al., 2015).

In summary, the FOCUS screen described in our study represents the first systematic dissection of functional OCT4-bound CREs which are essential for pluripotency maintenance. It illuminates a previously undescribed layer of regulatory mechanisms, one of which includes CREs and its target genes, in the overall pluripotency circuitry.

## ACKNOWLEDGEMENT

We thank Yu Tao, Fang Haitong, Nickolas Teo, Aloysi Aloysius Jun-Hui Quek and Samantha Seah for technical assistance and editorial suggestions. Y-H-L is supported by the [NRF Investigatorship Award – NRF12018-02], [JCO Development Programme Grant – 1534n00153)] and the [National Medical Research Council NMRC/CBRG/0092/2015].

## REFERENCE

Arnold, C.D., Gerlach, D., Stelzer, C., Boryn, L.M., Rath, M., and Stark, A. (2013). Genome-Wide Quantitative Enhancer Activity Maps Identified by STARR-seq. Science 339, 1074–1077.

Buecker, C., Srinivasan, R., Wu, Z., Calo, E., Acampora, D., Faial, T., Simeone, A., Tan, M., Swigut, T., and Wysocka, J. (2014). Reorganization of Enhancer Patterns in Transition from Naive to Primed Pluripotency. Cell Stem Cell 14, 838–853.

Bonev, B., Mendelson Cohen, N., Szabo, Q., Fritsch, L., Papadopoulos, G.L., Lubling, Y., Xu, X., Lv, X., Hugnot, J.-P., Tanay, A., et al. (2017). Multiscale 3D Genome Rewiring during Mouse Neural Development. Cell 171, 557–572.e24.

Boyer, L. A., Lee, T. I., Cole, M. F., Johnstone, S. E., Levine, S. S., Zucker, J. P., … & Gifford, D. K. (2005). Core transcriptional regulatory circuitry in human embryonic stem cells. Cell, 122(6), 947–956.

Chen, X., Xu, H., Yuan, P., Fang, F., Huss, M., Vega, V.B., Wong, E., Orlov, Y.L., Zhang, W., Jiang, J., et al. (2008). Integration of External Signaling Pathways with the Core Transcriptional Network in Embryonic Stem Cells. Cell 133, 1106–1117.

Cong, L., Ran, F.A., Cox, D., Lin, S., Barretto, R., Habib, N., Hsu, P.D., Wu, X., Jiang, W., Marraffini, L.A., et al. (2013). Multiplex Genome Engineering Using CRISPR/Cas Systems. Science 339, 819–823.

Dahéron, L., Opitz, S.L., Zaehres, H., Lensch, M.W., Lensch, W.M., Andrews, P.W., Itskovitz-Eldor, J., and Daley, G.Q. (2004). LIF/STAT3 signaling fails to maintain self-renewal of human embryonic stem cells. Stem Cells 22, 770–778.

Diao, Y., Fang, R., Li, B., Meng, Z., Yu, J., Qiu, Y., Lin, K.C., Huang, H., Liu, T., Marina, R.J., et al. (2017). A tiling-deletion-based genetic screen for cis-regulatory element identification in mammalian cells. Nature Methods 14, 629–635.

Diao, Y., Li, B., Meng, Z., Jung, I., Lee, A.Y., Dixon, J., Maliskova, L., Guan, K.-L., Shen, Y., and Ren, B. (2016). A new class of temporarily phenotypic enhancers identified by CRISPR/Cas9-mediated genetic screening. Genome Res. 26, 397–405.

Dixon, J.R., Selvaraj, S., Yue, F., Kim, A., Li, Y., Shen, Y., Hu, M., Liu, J.S., and Ren, B. (2012). Topological domains in mammalian genomes identified by analysis of chromatin interactions. Nature 485, 376–380.

Do, D.V., Ueda, J., Messerschmidt, D.M., Lorthongpanich, C., Zhou, Y., Feng, B., Guo, G., Lin, P.J., Hossain, M.Z., Zhang, W., et al. (2013). A genetic and developmental pathway from STAT3 to the OCT4-NANOG circuit is essential for maintenance of ICM lineages in vivo. Genes Dev. 27, 1378–1390.

ENCODE Project Consortium (2012). An integrated encyclopedia of DNA elements in the human genome. Nature 489, 57–74.

Fang, H.T., EL Farran, C.A., Xing, Q.R., Zhang, L.-F., Li, H., Lim, B., and Loh, Y.-H. (2018). Global H3.3 dynamic deposition defines its bimodal role in cell fate transition. Nature Communications 9, 1537.

Forristal, C.E., Wright, K.L., Hanley, N.A., Oreffo, R.O.C., and Houghton, F.D. (2010). Hypoxia inducible factors regulate pluripotency and proliferation in human embryonic stem cells cultured at reduced oxygen tensions. Reproduction 139, 85–97.

Fuellen, G., and Struckmann, S. (2010). Evolution of gene regulation of pluripotency--the case for wiki tracks at genome browsers. Biol. Direct 5, 67.

Fulco, C.P., Munschauer, M., Anyoha, R., Munson, G., Grossman, S.R., Perez, E.M., Kane, M., Cleary, B., Lander, E.S., and Engreitz, J.M. (2016). Systematic mapping of functional enhancer–promoter connections with CRISPR interference. Science 354, 769–773.

Grant, C.E., Bailey, T.L., and Noble, W.S. (2011). FIMO: scanning for occurrences of a given motif. Bioinformatics 27, 1017–1018.

Gu, P., Goodwin, B., Chung, A.C.K., Xu, X., Wheeler, D.A., Price, R.R., Galardi, C., Peng, L., Latour, A.M., Koller, B.H., et al. (2005). Orphan Nuclear Receptor LRH-1 Is Required To Maintain Oct4 Expression at the Epiblast Stage of Embryonic Development. Molecular and Cellular Biology 25, 3492–3505.

Kawazoe, S., Ikeda, N., Miki, K., Shibuya, M., Morikawa, K., Nakano, S., Oshimura, M., Hisatome, I., and Shirayoshi, Y. (2009). Extrinsic factors derived from mouse embryonal carcinoma cell lines maintain pluripotency of mouse embryonic stem cells through a novel signal pathway. Dev. Growth Differ. 51, 81–93.

Kearns, N.A., Pham, H., Tabak, B., Genga, R.M., Silverstein, N.J., Garber, M., and Maehr, R. (2015). Functional annotation of native enhancers with a Cas9–histone demethylase fusion. Nature Methods 12, 401–403.

Kim, H.-S., Tan, Y., Ma, W., Merkurjev, D., Destici, E., Ma, Q., Suter, T., Ohgi, K., Friedman, M., Skowronska-Krawczyk, D., et al. (2018). Pluripotency factors functionally premark cell-type-restricted enhancers in ES cells. Nature 556, 510–514.

Kobe, B., Kajava A. (2001) The leucine-rich repeat as a protein recognition motif. Current Opinion in Structural Biology 11.725–732.

Korkmaz, G., Lopes, R., Ugalde, A.P., Nevedomskaya, E., Han, R., Myacheva, K., Zwart, W., Elkon, R., and Agami, R. (2016). Functional genetic screens for enhancer elements in the human genome using CRISPR-Cas9. Nat. Biotechnol. 34, 192–198.

Lanner, F., and Rossant, J. (2010). The role of FGF/Erk signaling in pluripotent cells. Development 137, 3351–3360.

Li, D., Liu, J., Yang, X., Zhou, C., Guo, J., Wu, C., Qin, Y., Guo, L., He, J., Yu, S., et al. (2017). Chromatin Accessibility Dynamics during iPSC Reprogramming. Cell Stem Cell 21, 819–833.e6.

Lim, L.S., Loh, Y.-H., Zhang, W., Li, Y., Chen, X., Wang, Y., Bakre, M., Ng, H.-H., and Stanton, L.W. (2007). Zic3 is required for maintenance of pluripotency in embryonic stem cells. Molecular Biology of the Cell 18, 1348–1358.

Loh, Y.-H., Wu, Q., Chew, J.-L., Vega, V.B., Zhang, W., Chen, X., Bourque, G., George, J., Leong, B., Liu, J., et al. (2006). The Oct4 and Nanog transcription network regulates pluripotency in mouse embryonic stem cells. Nat. Genet. 38, 431– 440.

Ma, H., Ng, H.M., Teh, X., Li, H., Lee, Y.H., Chong, Y.M., Loh, Y.-H., Collins, J.J., Feng, B., Yang, H., et al. (2014). Zfp322a Regulates mouse ES cell pluripotency and enhances reprogramming efficiency. PLoS Genet. 10, e1004038.

Miyabayashi, T., Teo, J.-L., Yamamoto, M., McMillan, M., Nguyen, C., and Kahn, M. (2007). Wnt/beta-catenin/CBP signaling maintains long-term murine embryonic stem cell pluripotency. Proceedings of the National Academy of Sciences 104, 5668– 5673.

Onishi, K., and Zandstra, P.W. (2015). LIF signaling in stem cells and development. Development 142, 2230–2236.

Shen, Y., Yue, F., McCleary, D.F., Ye, Z., Edsall, L., Kuan, S., Wagner, U., Dixon, J., Lee, L., Lobanenkov, V.V., et al. (2012). A map of the cis-regulatory sequences in the mouse genome. Nature 488, 116–120.

Shlyueva, D., Stampfel, G., and Stark, A. (2014). Transcriptional enhancers: from properties to genome-wide predictions. Nat Rev Genet 15, 272–286.

Sokol, S.Y. (2011). Maintaining embryonic stem cell pluripotency with Wnt signaling. Development 138, 4341–4350.

Takahashi, K., and Yamanaka, S. (2006). Induction of Pluripotent Stem Cells from Mouse Embryonic and Adult Fibroblast Cultures by Defined Factors. Cell 126, 663– 676.

Zhang, J., Gao, Y., Yu, M., Wu, H., Ai, Z., Wu, Y., Liu, H., Du, J., Guo, Z., and Zhang, Y. (2015). Retinoic Acid Induces Embryonic Stem Cell Differentiation by Altering Both Encoding RNA and microRNA Expression. PLoS ONE 10, e0132566.

