## Supplemental Information for "Defining Essential Enhancer for Pluripotent stem cells using Features Oriented CRISPR-Cas9 Screen"

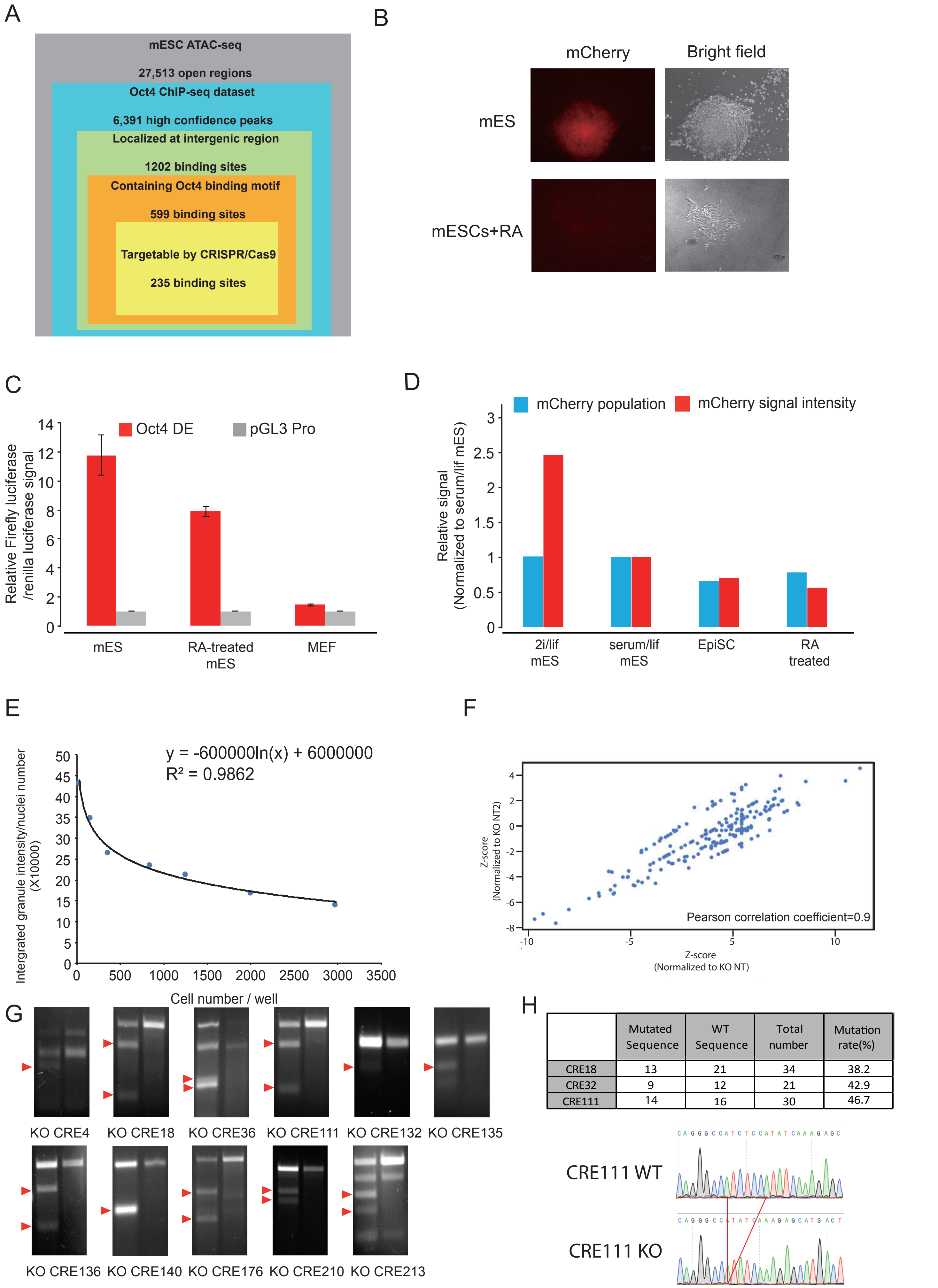


| | Mutated Sequence | WT Sequence | Total number | Mutation rate(%) |
|---|---|---|---|---|
| CRE18 | 13 | 21 | 34 | 38.2 |
| CRE32 | 9 | 12 | 21 | 42.9 |
| CRE111 | 14 | 16 | 30 | 46.7 |

Supplementary Figure 1

**Supplementary Figure 1. FOCUS identified important cis-regulatory elements for pluripotency maintenance**

A) Schematic of the pipeline used to generate FOCUS library targeting OCT4-bound cis-regulatory elements.
B) Immunofluorescence and brightfield images of untreated and RA-treated mESCs with Oct4 DE mCherry reporter (red). The mCherry reporter signal is lost upon treatment with RA.
C) Relative luciferase activity of Oct4 DE (red) in E14 cells, RA treated E14 cells and MEF. Data was normalized to pGL3 empty vector transfected cell (grey) which was set as 1. The bar chart shows mean ± SD of 2 independent experiments.
D) FACS result of Oct4 DE mCherry reporter E14 cells treated with different culture conditions. mCherry population percentage is shown in blue. mCherry signal intensity is shown in red.
E) Fitted curve of correlation between staining and cell number.
F) Scatter plot of Z-score normalized to KO NTs. Each dot represents one CRE. The x-axis is the Z-score of CREs obtained when normalized to NT1, while the y-axis indicates the Z-score of CREs obtained when normalized to NT2. The Pearson correlation coefficient is 0.9.
G) SURVEYOR assay confirmed specific targeting of CREs by sgRNAs. The red arrows indicate the expected band after SURVEYOR nuclease digestion.
H) Genotyping of sgRNA-transfected mESCs revealed 40% of editing efficiency of targeted *Oct4* motif.

**Related to Figure 1.**

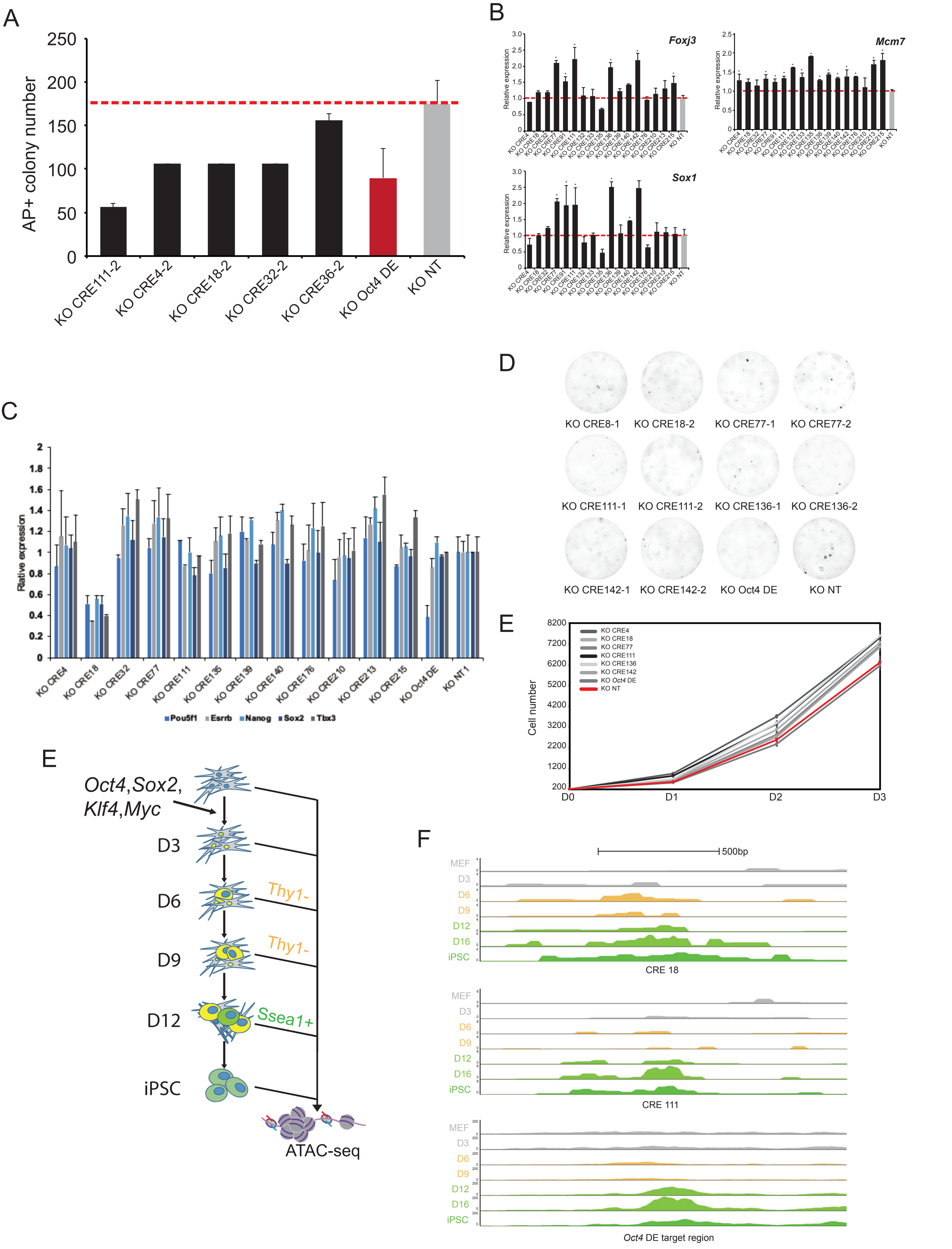


Supplementary Figure 2

**Supplementary Figure 2. Knockout of candidate CRE hits affects both the maintenance and establishment of pluripotent stem cells.**

A) Colony formation assay showing the knockout of selected CREs in D3 mESCs. The CREs are shown in black, the negative control is shown in grey and the positive control, *Oct4* DE, in red. The bar chart shows mean ± SD of 3 replicates.
B) qRT-PCR showing increased expression of differentiation genes when hit CREs were knocked out during RA-induced differentiation. The CREs are shown in black, while the control knock out is in grey. The bar chart shows mean ± SD of 3 independent experiments. Student’s t-test was used for statistical analysis. * represents p-value <0.05.
C) Images of AP staining of mouse somatic cell reprogramming at Day 12 post OSKM induction.
D) Cell proliferation curve of MEFs with CREs knocked out using CRISPR. Graph shows mean ± SD of 3 replicates.
E) Schematic for the ATAC-seq libraries from different time points of somatic cell reprogramming.
F) UCSC screenshot of ATAC-seq signal of candidate CREs at different time points of mouse somatic cell reprogramming.

**Related to Figure 2.**

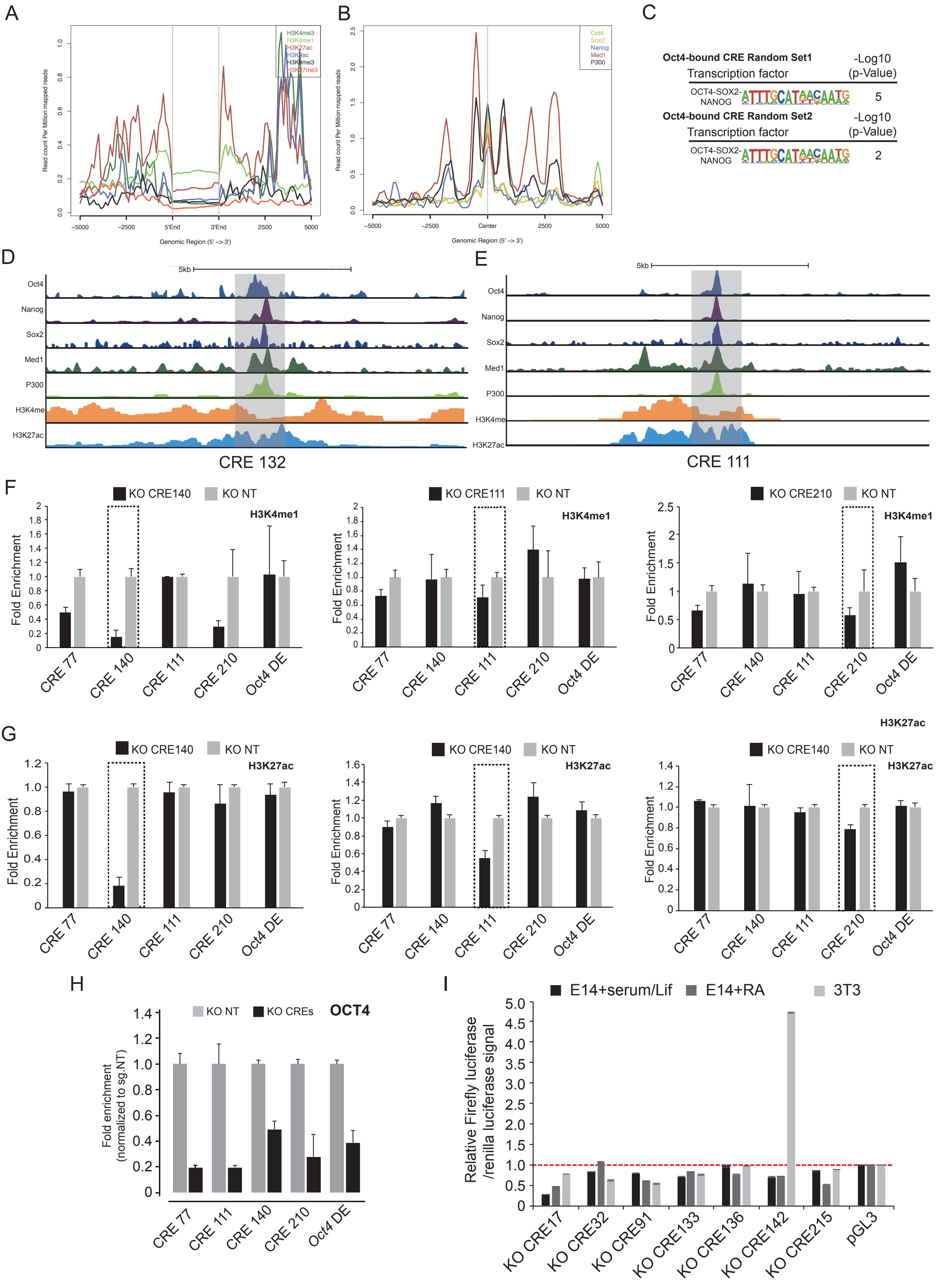


Supplementary Figure 3

**Supplementary Figure 3. Hit CREs displayed enhancer characteristics and activity in mESCs.**

A) Average enrichment plot of histone marks on non-hit CREs. Low enrichment of active enhancer markers, H3K4me and H3K27ac, was observed. Different histone marks are indicated as in different colours.
B) Average enrichment plot of transcription factors binding on non-hit CREs. Low enrichment of transcription factors was observed on non-hit CREs. Transcription factors are indicated in different colours.
C) Motifs enriched at non-hit Oct4-bound intergenic CREs sites.
D) UCSC browser view of enriched histone marks and transcription factors at CRE 132. 1.5 kb enhancer region is highlighted in grey.
E) UCSC browser view of enriched histone marks and transcription factors at CRE 111. 1.5 kb enhancer region is highlighted in grey.
F) ChIP-qPCR results for the knockout of candidate CREs. Decreased enrichment of H3K4me1 was observed. Black bars represent CRE KO group. Grey bars represent KO NT control. The bar chart shows mean ± SD of 2 independent experiments.
G) ChIP-qPCR analysis showed that knockout of candidate CREs reduced the enrichment of H3K27ac. Black bars represent CRE KO group. Grey bars represent KO NT control. The bar chart shows mean ± SD of 2 independent experiments.
H) Knockout of the candidate CREs (black) reduced the enrichment of OCT4 binding on-site. ChIP-qPCR enrichment for each CREs was normalized to KO NT (grey) and presented as fold enrichment. Data are representative of at least 3 independent experiments. The bar chart shows mean ± SD.
I) Relative luciferase activity of reporter plasmids (pGL3-promoter) containing a fragment of candidate CREs in E14 (black), RA treated E14 cells (dark grey) and 3T3 (light grey). The bar chart shows mean ± SD of 3 replicates.

**Related to Figure 3.**

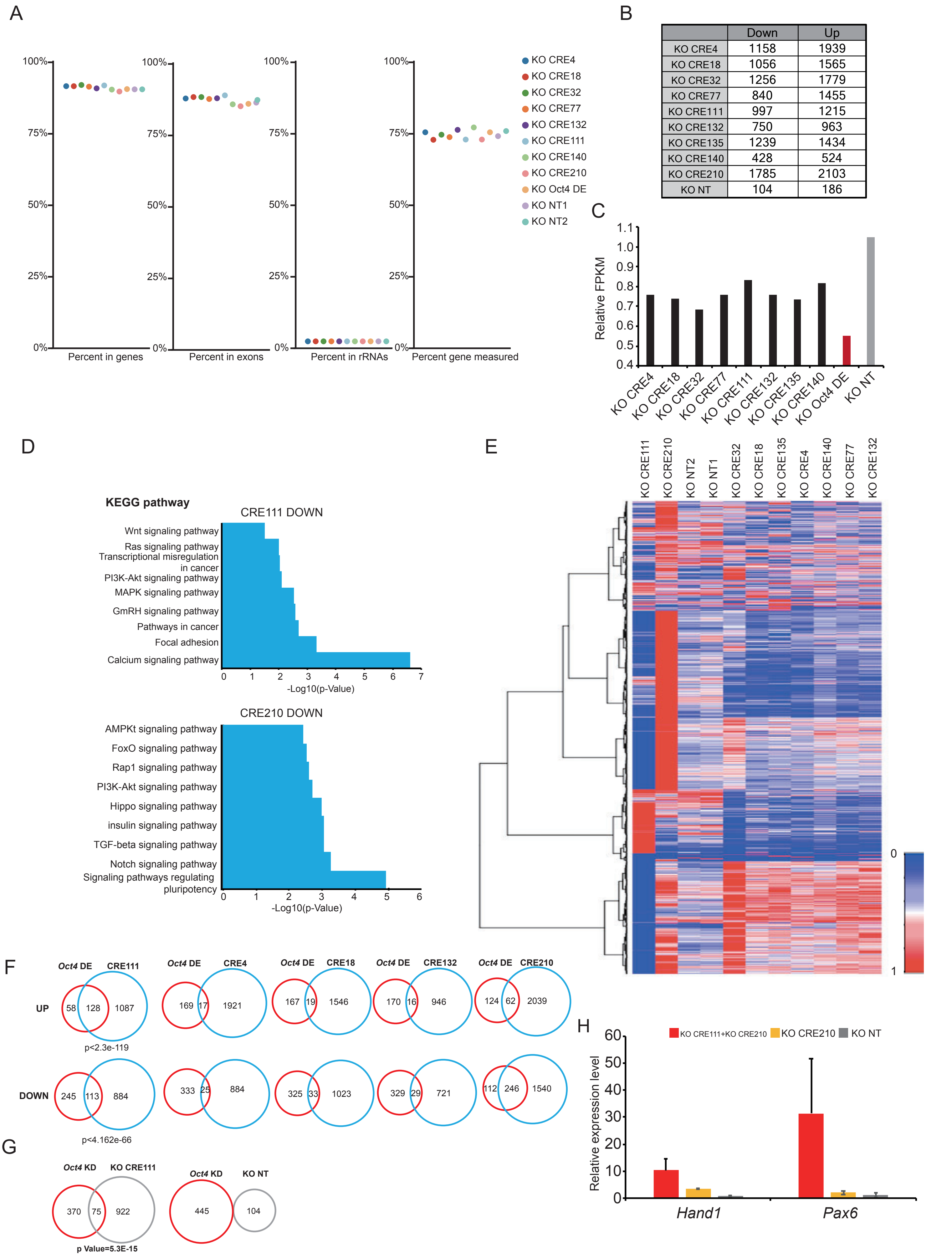


### Supplementary Figure 4

**Supplementary Figure 4. Transcriptomic profile elicited by CRE KO.**

A) QC plot for KO CRE RNA-seq libraries.
B) The number of up- and down-regulated genes for each KO CRE RNA-seq library.
C) Relative FPKM for *Oct4* expression in each KO CRE RNA-seq library. Black bars represent candidate CREs. Red bar represents *Oct4* DE, which was used as positive control. Grey bar represents KO NT control.
D) Heatmap showing the clusters of all KO CRE RNA-seq libraries.
E) KEGG pathway analysis of down-regulated genes in KO CRE111 and KO CRE 210 RNA-seq libraries. X-axis represents the –Log10(p-Value). Y-axis represents GO term.
F) Venn diagram showing the number of overlapped genes which were differentially expressed between CRE KO and Oct4 DE KO. p-values were calculated to test the significant level of the overlap.
G) Venn diagram showing the number of overlapping differentially expressed genes between CRE111 and *Oct4* knockdown. p-value was calculated to test the sigificance of the overlap.
H) Combinatorial KO of CRE111 and CRE210 further increased the expression of differentiation genes, *Hand1* and *Pax6*, compared to single KO. Cells transfected with KO NT sgRNA was used as control (grey) and all data was normalized to it. The bar chart shows mean ± SD of 3 replicates.
I)

**Related to Figure 4.**

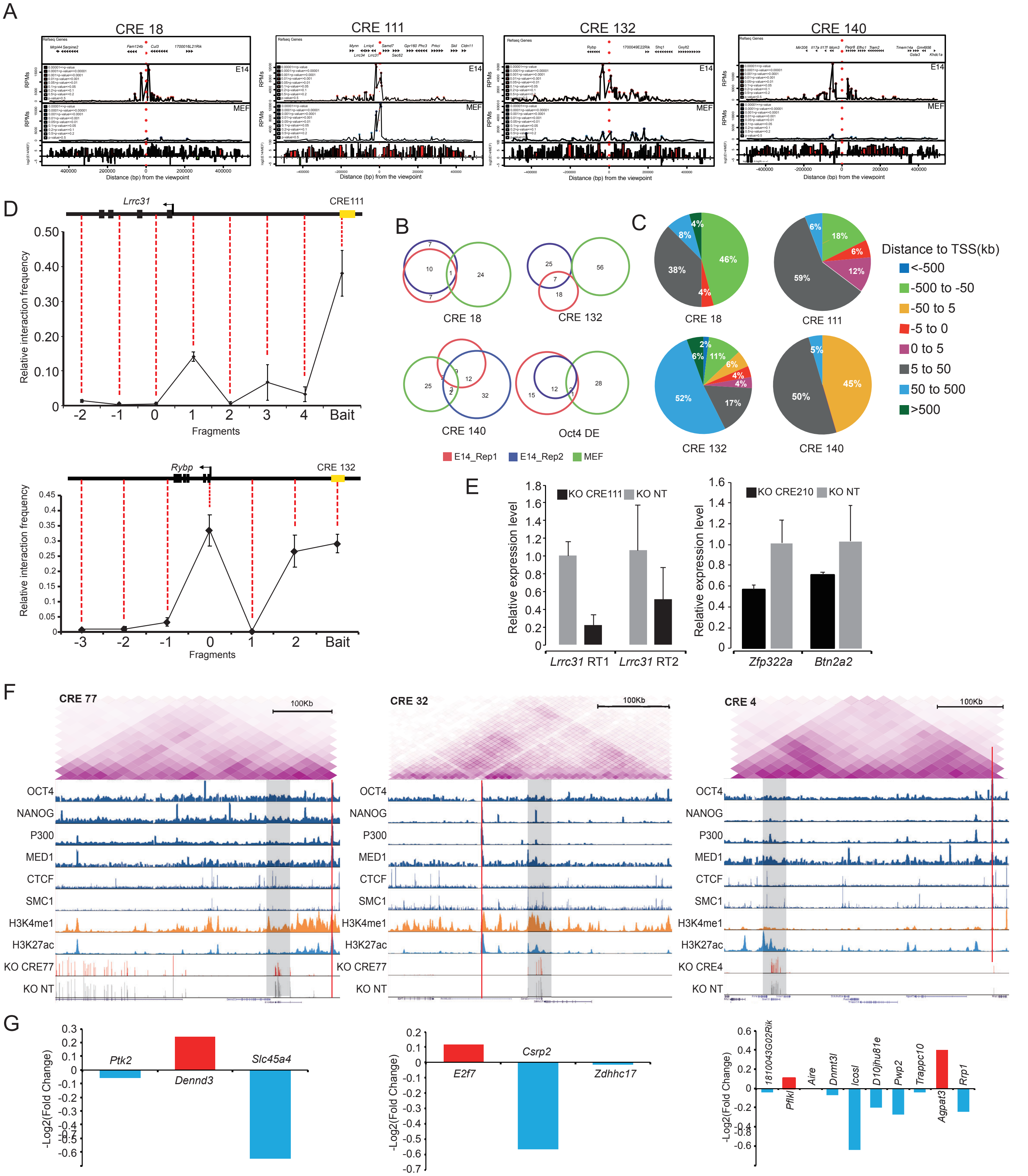


Supplementary Figure 5

**Supplementary Figure 5. Candidate CREs interact in 3D genomic space with novel and known pluripotency genes to regulate pluripotency.**

A) Interaction of CREs and its target loci was detected in E14 cells using 4C-seq. Interactions ($q<0.05$, r3Cseq) in E14 cells (2nd panel) and MEFs (3rd panel) are shown. The 4C signal plot (in units of RPM) was compared between E14 and MEF and plotted using r3Cseq package (4th panel).
B) Overlap of significant interacting regions of hit CREs between biological replicates of E14 (red and blue) and MEF (green).
C) Pie chart showing the distance of CRE interacting loci to their nearest TSSs.
D) The interaction of CRE111 (top) and CRE132 (bottom) to its target gene in E14 cells was validated using 3C. X-axis shows 3C fragment corresponding to actual genomic location. Y-axis shows relative interacting frequency between E14 and BAC control. The corresponding location of each fragment in the genome is indicated using the red dotted lines. The bar chart shows mean ± SD of 3 replicates.
E) qRT-PCR showing decreased putative targets' expression in CRE KO, which is shown in black; KO NT control is shown in grey. The expression is normalized to KO NT controls. The bar chart shows mean ± SD of 3 replicates.
F) Integrative genomic view of Hi-C, ChIP-seq and RNA-seq data of CRE 77 (left), CRE32 (middle) and CRE4 (right). Top panel: Hi-C interactions within the same TAD of CRE. Middle panel: ChIP-seq UCSC browser of pluripotency and architectural proteins, as well as active enhancer marks. Bottom panel: RNA-seq UCSC browser of KO CRE and KO NT. The location of CRE is highlighted using red line. Putative target of CRE is indicated using grey shades.
G) The bar chart showing the expression change of genes located within the same TAD as CRE77 (left), CRE32 (middle) and CRE4 (right) when CRE was knocked out. The y-axis represents the -Log2(Fold change). The genes with decreased expression are shown as the blue bars. The genes with increased expression are shown as the red bars.

**Related to Figure 5.**

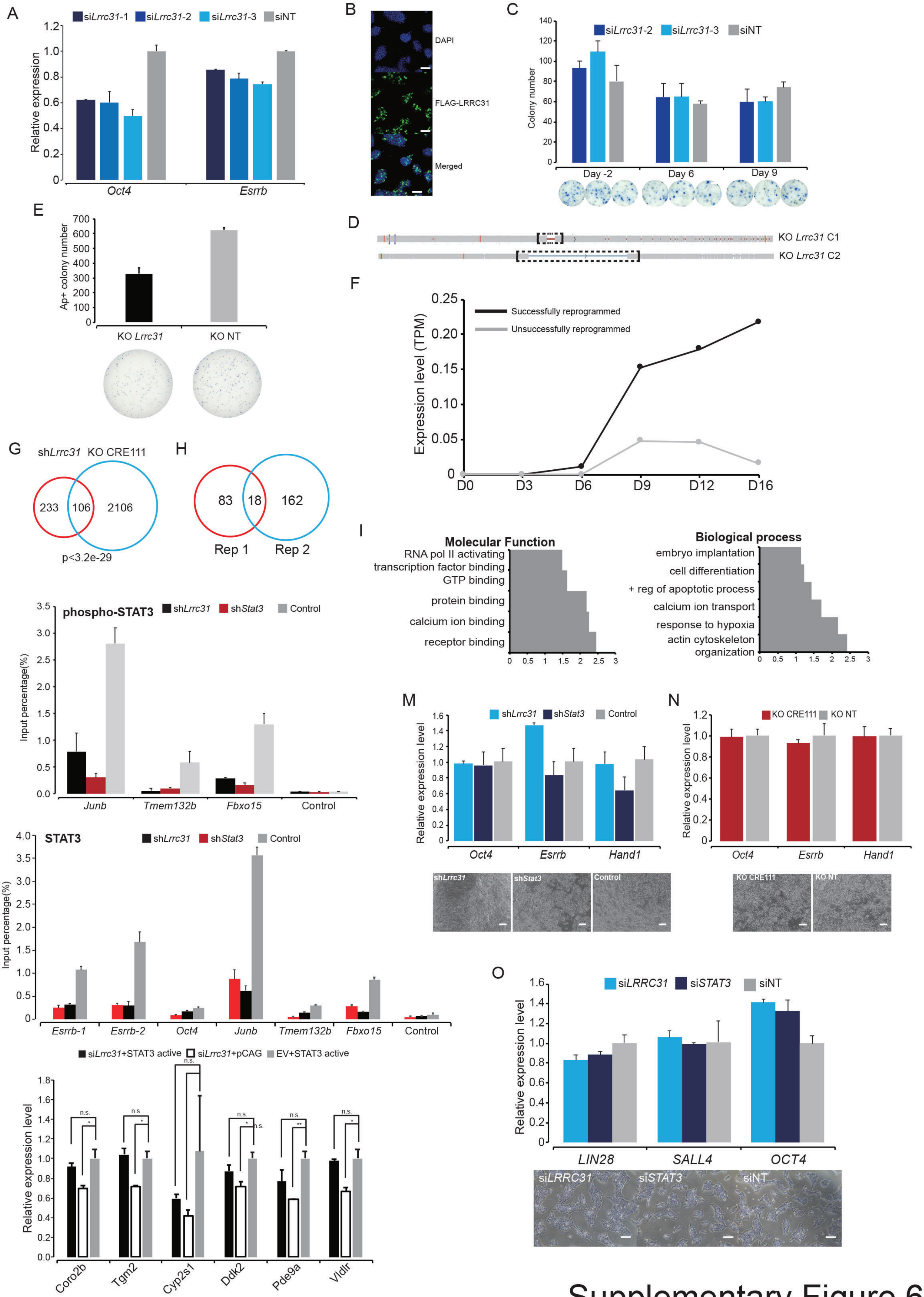


Supplementary Figure 6

**Supplementary Figure 6. *Lrrc31*, a target gene of CRE111 regulates the maintenance of pluripotency in mES cells.**

A) qRT-PCR showing the knockdown of *Lrrc31* using siRNA, which resulted in decreased expression of *Oct4* and *Esrrb*. *Lrrc31* siRNAs are shown in shades of blue while siNT control is shown in grey. The bar chart shows mean ± SD of 3 replicates.
B) Immunofluorescence images of FLAG-LRRC31. Staining was performed with anti-FLAG antibody (green). Nuclei were stained with Hoechst stain (blue). Scale bar is 200 μm.
C) Knocking down of *Lrrc31* at different time points of somatic cell reprogramming elicited varied phenotypes. Top panel: reprogramming efficiency (AP+ colony numbers) was assayed at 12 d.p.i. *Lrrc31* knockdown (blue and dark blue) at 9 d.p.i decreased reprogramming efficiency compared to siNT (grey). The bar chart shows mean ± SD of 3 replicates. Bottom panel: Representative images of each treatment group.
D) Sequencing result confirmed the truncated *Lrrc31* coding sequencing in *Lrrc31* KO C1 and C2.
E) Colony formation assay showing that the knock-out of *Lrrc31* compromised self-renewal of mESCs. The bar chart shows mean ± SD of 3 biological replicates. Representative images are shown below the bar chart.
F) Expression profile of *Lrrc31* in successfully and unsuccessfully reprogrammed cells. Y-axis is expression level (TPM). The x-axis represents different time points throughout the reprogramming process.
G) Venn diagram showing the overlap between *Lrrc31* KD RNA-seq and KO CRE 111 RNA-seq. p-values were calculated to test the significant level of the overlap.
H) GO analysis of differentially expressed genes upon *Lrrc31* knockdown. X-axis represents the –Log10(p-Value). Y-axis represents GO terms.
I) Venn diagram showing the overlap of identified proteins between two biological replicates of FLAG-LRRC31 pull-down assay.
J) ChIP qRT-PCR demonstrating that knocking down of *Lrrc31* decreased the binding of phospho-STAT3 on its targets. Samples treated with shRNA against *Lrrc31* are shown as black. pSUPER Empty vector was used as control (light grey). The shRNA of *Stat3* was used as positive control (dark red). The bar chart shows mean ± SD of 3 replicates.
K) ChIP qRT-PCR demonstrating that knocking down of *Lrrc31* decreased the binding of STAT3 on its target. The sample treated with shRNA against *Lrrc31* is shown as a black bar. pSUPER Empty vector was used as control (light grey). The shRNA of *Stat3* was used as positive control (dark red). The bar chart shows mean ± SD of 3 replicates.
L) Rescue of differentially expressed genes of *Lrrc31* knockdown by STAT3 dominant active mutant overexpression in Lrrc31 knockdown ES-E14. The bar chart shows mean ± SD of 3 replicates. Student's t-test was used for statistical analysis. * represents p-value <0.05.

M) qRT-PCR showing the knockdown of *Lrrc31* in F9 cells resulted in no significant change in the expression of marker genes. The bar chart shows mean ± SD of 2 independent experiments. Corresponding brightfield images for each treatment group is shown below. Scale bar is 200 µm.

N) qRT-PCR showing that the knockout of CRE111 gave rise to insignificant change in the expression of marker genes. The bar chart shows mean ± SD of 2 independent experiments. The corresponding brightfield images for each treatment group is shown below. Scale bar is 200 µm.

O) qRT-PCR showing that the knockdown of *LRRC31* showed no significant change in the expression of marker genes in hESCs. The bar chart shows mean ± SD of 2 independent experiments. The corresponding brightfield image for each treatment group is shown below. Scale bar is 200 µm.

**Related to Figure 6.**

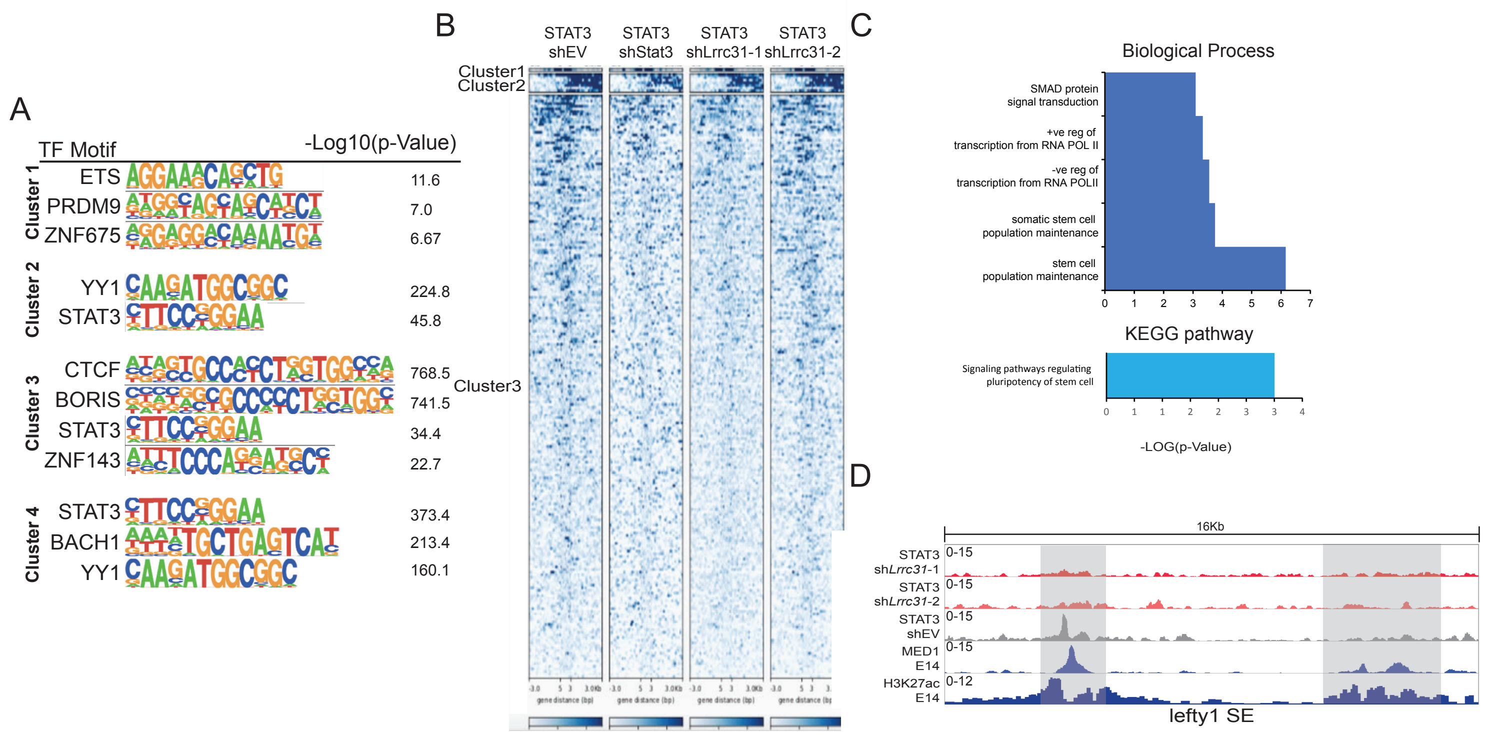


Supplementary Figure 7

**Supplementary Figure 7. *Lrrc31* regulated the binding of STAT3 to the super-enhancers.**

A) Motifs enriched at STAT3 binding sites of different clusters.
B) Enrichment of STAT3 on super enhancers of mESCs in the indicated conditions.
C) Gene ontology analysis of STAT3-bound super enhancers' target genes. X-axis represents the –Log10(p-Value) and Y-axis represents GO term.
D) UCSC screenshot of binding profile of STAT3 on *Lefty1* super enhancer (indicated by the binding of MED1 and H3K27ac) when *Stat3* or *Lrrc31* was knocked down.

**Related to Figure 7.**

#### EXPERIMENT PROCEDURE

##### sgRNA design

The *Oct4* motif were identified in CREs where only one strong OCT4 binding site was detected by the published OCT4 ChIP-seq (Chen et al, 2008). If more than one *Oct4* motif was detected, the one which is nearest to the actual OCT4 binding site was chosen as the target motif. Then the sequence of all OCT4 binding site with *Oct4* motif was imported into CRISPR sgRNA designer (Doench et al., 2016). Only the sgRNAs cut within the motif of the binding site were chosen for downstream screen. For batch 2 sgRNAs, sgRNA Scorer (Mali et al., 2015)were used to design alternative sgRNA cutting within *Oct4* motif or maximum 20bp around the motif sequence.

##### Sg.CRISPR Screen

96-well plates were coated with 0.1% gelation overnight at 37℃ incubator. 5K E14 cells was seeded onto each well of 96-well plate. Transfected was conducted 24 hours post seeding. 0.3ul of lipo2000, 100ng of plasmids DNA and 9ul opti-MEM was mixed and incubated at room temperature for 20min before adding into each well of 96 well plates. Cell was selected using puromycin at a concentration of 1ug/ml from D1 to D4 post transfection. Cell was fixed with 4% PFA for 15 min at room temperature. After blocking with 7% FBS for 30 mins at room temperature, cell was stained with Oct4 antibody (1:1000) overnight at 4℃. Cell was washed with PBS for three times at room temperature. And the cell was stained with Goat anti-Rabbit IgG (H+L), Alexa Fluor® 488 at room temperature for 2 hours. The cell was stained with Hoechst(1ug/ml) for 15min at room temperature after being washed for three times using PBS. The cell was washed for another three times in PBS and covered in 100ul PBS.

Plates were imaged using IXU ultra plate-scanning confocal microscope (molecular device) at 20X magnification. For each well of 96-well plate, both OCT4 immunofluorescence and DAPI image were taken in the 25 fields of the well to represent each well. Then the average Oct4 immunofluorescence for each nuclei (identified by DAPI) was calculated by MXexpress software for each well. To normalize such bias in our screen result, the correlation between the cell number and Oct4 staining intensity were generated. The average OCT4 immunofluorescence was normalized using the regression formula. Z-score were calculated as previously described(Toh et al., 2016). A cutoff at -2 was used to identify hits from the CRISPR screen.

**Colony formation assay**

E14 was transfected with LentiCRISPR plasmids using lipo 2000. Cell was selected with puromycin(1ug/ml) from Day 1 post transfection to Day 3. Then cell was counted and 2K cells was seeded into one well of 12 well plates for each binding site KO. Cell was fixed with 4% PFA and stained with VECTOR Blue Alkaline Phosphatase (Blue AP) Substrate Kit following the manufacturer's protocol. Colony number for each well was automated counted using CellProfiler (Carpenter et al., 2006).

**Luciferase assay**

The potential enhancers from the primary was PCR amplified and cloned into KpnI and XhoI site in pGL3-promoter vector. Both E14 and 3T3 were co-transfected with pGL3-promoter vector and pRL renilla luciferase control vector using lipo2000. Cell was harvested with passive lysis buffer 72 hours post transfection. Then luciferase assay was performed according to manufacturer's protocol using GloMAX Explorer system.

**SUVEYOR assay**

E14 was transfected with LentiCRISPR plasmids using lipo 2000 as described above. Cell was selected with puromycin (1ug/ml) from Day 1 post transfection to Day 3. Genomic DNA was extracted using QIAGEN DNeasy Blood & Tissue Kit following manufacturer's protocol.

Genomic DNA surrounding the CRISRP target site for each gene was PCR amplified and purified using QIAquick PCR Purification Kit following manufacturer's protocol. 400ng of the purified PCR products were mixed with 2μl 10X Taq polymerase PCR buffer (QIAGEN) and nuclease free water to a final volume of 20μl. PCR product mixture was then re-annealed: 95℃ for 10min, 95℃to 25℃ ramping at – 0.25℃/s, and 25℃ forever. After re-annealing, products were treated with SURVEYOR nuclease and SURVEYOR enhancer S (IDT), following the manufacturer's recommended protocol, and analyzed on 2% agarose gels with Ethidium bromide. Gel was imaged with Gel Doc gel imaging system (Bio-rad).

**mESC culture**

E14 and D3 mouse ES cells were cultured on gelatin (0.1%) coated plates in mESC medium at 37°C with 5% CO2. The medium composition: 15% heat-inactivated

ES-qualified fetal bovine serum (GIBCO), 2mM L-glutamine (GIBCO), 0.1mM nonessential amino acid (NEAA, GIBCO), 5000U/ml penicillin/streptomycin (GIBCO), 0.055mM β-mercaptoethanol (GIBCO) and 1000U/ml home-made leukemia inhibitory factor in Dulbecco's modified Eagles' medium (DMEM, Hyclone).

**F9 culture**

F9 mouse embryonal carcinoma cells were cultured on gelatin (0.1%) coated plates in mESC medium at 37°C with 5% CO2. The medium composition: 10% heat-inactivated fetal bovine serum (GIBCO), 2mM L-glutamine (GIBCO), 0.1mM nonessential amino acid (NEAA, GIBCO), 5000U/ml penicillin/streptomycin (GIBCO) in Dulbecco's modified Eagles' medium (DMEM, Hyclone).

**hESC culture**

The human embryonic stem cells line H1 (WA-01) were cultured on Matrige l(Corning) coated plates in mTeSR (Stem cell technology) medium at 37°C with 5% CO2. H1 was passaged every 4-5 days using using L7 hPSC passage solution according to manufacturer's protocol and replated at a ratio of 1:10.

**Reprogramming**

Immortalized MEF(iMEF) was infected with Lentivirus contain both sgRNA and Cas9. iMEF were selected with puromycin at 3.3ug/ml for 3 days. Then 2K iMEF were seeded onto one well of gelated coated 12-well plates and infected with OSKM virus simultaneously. Reprogramming medium were changed everyday. Cell was fixed with 4% PFA and stained with VECTOR Blue Alkaline Phosphatase (Blue AP)

Substrate Kit following the manufacturer's protocol at Day 10 post infection. Colony number for each well was automated counted using CellProfiler.

**shRNA transfection in mESC**

Dharmacon siDESIGN center was used for the design of the shRNA sequences (http://dharmacon.gelifesciences.com/design-center/). shRNA sequences were ordered as DNA oligos from IDT and cloned into pSUPER-puro plasmid. mESCs were trypsinized for 5 minutes at 37 °C, counted by hemocytometer and 200,000 cells seeded into one well of 6 well plates. mESCs were transfected with 2.5 µg of shRNA construct and 5 µl of Lipofectamine 2000 (Thermofisher). 1 µg/ml of puromycin was added to the mES medium 24 hours after transfection. All the knockdown experiments were carried out in biological triplicates.

**siRNA transfection in mESC and hESC**

siRNA was reversed transfected into mESC and hESC at a final concentration of 25nM using DharmaFECT 1 reagent (Dharmacon). RNA was harvested 72hrs post reverse transfection for downstream assay.

**ChIP**

ChIP was performed as described previously (Loh et al., 2006). Briefly, ten million cells were cross-linked with 1% formaldehyde for 10 minutes at room temperature; formaldehyde was then quenched by the addition of 200 mM glycine and cells were washed once with ice-cold PBS.

Cell lysis was carried out with a lysis buffer containing 10 mM Tris-Cl (pH 8), 100 mM NaCl, 10 mM EDTA, 0.25% Triton X-100 and protease inhibitor cocktail

(Roche). Cells were then resuspended in 1% SDS lysis buffer containing 50 mM HEPES-KOH (pH 7.5), 150 mM NaCl, 1% SDS, 2 mM EDTA, 1% Triton X-100, 0.1% NaDOC and protease inhibitor cocktail. The suspension was nutated for 15 minutes in the cold room before spinning down to collect the chromatin pellet. The pellet was then washed two times with 0.1% SDS lysis buffer containing 50 mM HEPES-KOH (pH 7.5), 150 mM NaCl, 0.1% SDS, 2 mM EDTA, 1% Triton X-100, 0.1% NaDOC, 1 mM PMSF and protease inhibitor cocktail. Sonication was conducted on the ice, using the Branson Sonifier 250 (11 cycles, power amplitude of 35%, 30 sec pulses on with 59.9 s pulses off). The chromatin solution was clarified by centrifugation at 20,000 g at 4°C for 45 minutes and then pre-cleared with Dynabeads protein A (Life Technologies) for 2 hours at 4°C. The pre-cleared chromatin sample was incubated with 50 μl of Dynabeads protein A loaded with 5 μg antibody overnight at 4°C. The beads were washed three times with 0.1% SDS lysis buffer, once with 0.1% SDS lysis buffer/0.35 M NaCl, once with 10 mM Tris-Cl (pH 8)/1 mM EDTA/0.5% NP40/0.25% LiCl/0.5% NaDOC, and once with TE buffer (pH8.0). The immunoprecipitated material was eluted from the beads by heating for 45 minutes at 68°C in 50 mM Tris-Cl (pH 7.5), 10 mM EDTA, 1% SDS. To reverse the crosslinks, samples were incubated with 1.5 μg/ml Pronase at 42°C for 2 hr followed by 67°C for 6 hr. The samples were then extracted with phenol-chloroform-isoamyl alcohol (25:24:1) followed by chloroform, ethanol precipitated in the presence of glycogen, and re-suspended in TE buffer.

**Co-IP/MS**

E14 cells was transfected with LRRC31-FLAG-Puro overexpression plasmid using Lipo2000 (Thermo Fisher) according to manufacturer’s protocol. Cell was then selected with puromycin at 1 μg/ml for 96 hrs. 10 million LRRC31-FLAG transfected E14 cells were harvested and lysed with IPH buffer (50 mM Tris-HCl, pH 8.0, 150 mM NaCl, 5 mM EDTA, 0.5% NP-40, with Roche's cocktail protease inhibitors). IP was carried out using EZviewTM red anti-FLAG M2 affinity beads overnight at 4°C. After washing with IPH buffer three times, beads were re-suspended in SDS loading buffer, boiled for 5min, followed by SDS-PAGE and mass spectrometry.

**Immunoflourescence (IF)**

Cell were fixed with 4% paraformaldehyde for 20 min, permeabilized with 0.2% Triton X-100 for 10 min, and blocked with 7% FBS in PBS for 30 min at room temperature. After blocking, cells were stained with anti-FLAG antibody (Sigma, F1804) followed by Alexa 488 conjugated secondary antibody (Thermo Fisher) and counterstained with Hoechst 33342 (Thermo Fisher). Images were captured with Zeiss LSM700 confocal microscope.

**Circularized Chromosome Conformation Capture (4C)-seq**

4C library were prepared using previously published protocol(Stadhouders et al., 2013). In summary, 10 million E14 cells and MEF were lysed with cross-linked by 1% formaldehyde for 10 min at room temperature. Cross-linked chromatin was digested with the primary restriction enzyme HindIII (CRE18, CRE111, CRE132, Oct4 DE; NEB), EcoRI(CRE 140; NEB); Next, 3C library was generated by proximity ligation using T4 DNA ligase (EL0013, Thermo Scientific) on HindIII digested DNA, followed by cross-link removal using Proteinase K (AM2546, Vivantis). The 3C libraries were then subjected to a second restriction enzyme digestion using DpnII (CRE132, Oct4 DE; NEB), NlaIII(CRE18; NEB), CviQI(CRE111, CRE140; NEB) followed by another ligation reaction using T4 DNA ligase. For each viewpoint, 3.2 μg of the resulting 4C templates was used to perform an inverse PCR. The PCR products were then run on 2% agarose gels. On the gel, smears from 200 to 600 bp were excised and unwanted PCR product bands were removed. DNA was then extracted from the cut-out gel pieces for next-generation sequencing on an Illumina Miseq. Interaction regions for each bait sequence was identified using r3Cseq package(Thongjuea et al., 2013).

**Chromosome conformation capture (3C)**

Ten million cells were harvested and washed twice with ice-cold PBS. Cells were then re-suspended in 12 ml PBS containing 10% fetal bovine serum. 0.649 ml of 37% formaldehyde (Sigma) was added to the mixture, and the mixture was nutated for 10 minutes at room temperature. The reaction was quenched with 1.6 ml of cold 1 M glycine, followed by washing twice with 14 ml ice-cold PBS. The pellet for each cell line was re-suspended in 5 ml of lysis buffer [10 nM Tris–HCl, pH8, 10 mM NaCl, 0.2% NP-40 and 1X Protease inhibitor] and incubated at 4°C for 10 minutes. Nuclei were then pelleted through centrifuging and resuspended in 500 μl of NEB Buffer 3.1 (New England BioLabs). 7.5 μl of 20% SDS was added to each tube, and mixed for 1 h at 37°C, at 900 r.p.m. on a thermomixer (Eppendorf). This was followed by the addition of 50 μl of 20% Triton-X100 (Sigma-Aldrich), and further mixing for 1 h at

37°C, at 900 r.p.m to quench SDS. 400U of the BglII enzyme (NEB) was added to each tube, followed by overnight incubation at 37°C and 900 r.p.m.

A 5 µl aliquot of digested DNA was assessed for completeness of digestion using qRT-PCR. The BglII enzyme was inactivated by the addition of 40 µl of 20% SDS and mixing for 25 minutes at 65°C at 900 r.p.m. The incubated samples were added to 6.125 ml of 1.15× ligation buffer (NEB) and 375 µl of 20% Triton-X100, with mixing for 1 h at 37°C. The cross-linked digested DNA was re-ligated by the addition of 100 U T4 DNA ligase (Thermofisher) and an overnight incubation at 16°C. 15 µl of 20 mg $ml^{-1}$ Proteinase K (Promega) was then added to the mixture, which was incubated at 65°C overnight.

The samples were cooled to room temperature, and 30 µl of 10 mg $ml^{-1}$RNAse A (QIAGEN) was added and the samples were incubated at 37°C for 1 hour. Then the 3C library was extracted by phenol-chloroform extraction and purified using PCR purification kit (QIAGEN). The 3C interactions were detected by qPCR and 2× SYBR Green Mastermix (KAPA) following the qRT-PCR protocol.

**qRT-PCR**

Total RNA was extracted using the Trizol reagent (Invitrogen). Contaminating DNA was removed by DNaseI (Ambion) treatment, and the RNA was further purified using QIAGEN RNeasy Kit. First strand cDNA was synthesized using the iScript cDNA synthesis kit (Bio-Rad) according to the manufacturer’s instructions. Quantitative real-time PCR was performed on the CXF384 Real-time System (Bio-Rad), using a Kapa SYBR Fast qPCR kit (Kapa Biosystems). The expression level of each gene was normalized to the expression level of *Gapdh*.

**RNA-seq preparation**

Cellular RNA was extracted using the RNA extraction protocol mentioned in previous section. The quality of extracted RNA samples was accessed by qRT-PCR. mRNA libraries were prepared using the TruSeq Stranded mRNA Library Prep Kit (Illumina, RS-122-2101), following the manufacturer's protocol. In brief, mRNA was purified from total RNA by incubating with poly-T oligo-attached magnetic beads. Following purification, mRNA was fragmented into small pieces using divalent cations at an elevated temperature. The cleaved RNA fragments were reverse transcribed into first strand cDNA using reverse transcriptase and random primers. The second strand cDNA was then synthesized using DNA Polymerase I and RNase H. These strand-specific cDNA fragments undergo an end repair process to produce a single 'A' base overhang, followed by ligation of the adapters. The products were purified and enriched with PCR to create the final cDNA library. Finally, samples were pooled and sequenced using Hiseq 2000 platform with a read length of 100bp.

**Western blot**

Cells were lysed with IPH buffer supplemented with protease inhibitor cocktail and PMSF. 50 μg of cell lysate was loaded onto an SDS-PAGE gel and transferred onto a polyvinylidene difluoride (PVDF) membrane (Bio-Rad). The membrane was blocked with 5% skim milk at room temperature for 1 hour, followed by overnight incubation with anti-phospho-STAT3(abcam, ab76315), anti-STAT3(abcam, ab119352) and anti-ACTIN (Santa Cruz) antibodies at 4°C. The blot was subsequently incubated with horse-radish peroxidase (HRP)-conjugated anti-mouse IgG (1:10,000, Beith). The HRP signals were detected using SuperSignal West Dura Extended Duration Substrate (Thermofisher) and captured using Gel Doc system (Bio-rad).

**CRE KO during early embryo development**

crRNA targeting at candidate CRE was synthesized by IDT. crRNA and tracrRNA were resuspended using injection buffer(1mM Tris-HCl, pH7.5; 0.1mM EDTA) to a final concentration of 1μg/μl. Then 5 μg of crRNA was mixed with 10 μg of tracrRNA and annealed in a thermocycler (95°C, 5 min; ramp down to 25°C at 5°C/min). Then the cr/tracrRNA mixture was mixed with Alt-R S.p. Cas9 Nuclease 3NLS (IDT) to a final concentration of 20 ng/μl for both guide RNA and Cas9 protein, followed by 15 min incubation at room temperature. The injection mix was then centrifuged at 13,000 rpm for 5 min at room temperature and pass through a Millpore filter(UFC30VV25). The microinjection was performed following published microinjection procedure(Harms et al., 2001).

**Average enrichment plot**

ATAC-Seq libraries(Fang et al., 2018) for mouse reprogramming cells were mapped to mm9 genome using STAR aligner (cite) allowing upto 3 mismatches. The options –alignIntronMax and –alignEndsType were set to 1 and EndToEnd respectively. The mapped bam files were then used as an input for ngsplot (cite) to determine the average enrichment of these libraries over the identified enhancers.

**ChIP-Seq Analysis**

ChIP-Seq libraries (Histones from ENCODE, Transcription Factors were from (Chen et al., 2008)) were mapped in a similar way to the ATAC-Seq libraries in order to determine their average enrichment over the identified enhancers. Moreover, the makeTagDirectory and makeUCSCfile scripts of HOMER (Heinz et al., 2010) were used to generate the UCSC screenshots. Peaks of Oct4 ChIP-Seq library were determined using the findPeaks script of HOMER with default settings for transcription factors.

**Motif enrichment**

findMotifsGenome.pl script of HOMER was used in order to determine the motifs found in the identified enhancers.

**Determination of CREs**

MACS2 (Zhang et al., 2008) was used to determine accessible sites in the mES ATAC-Seq library(Li et al., 2017) using the following parameters: "-g mm --outdir ./MACS2/2i/ --nolambda --nomodel --keep-dup all –call-summits". These site were then correlated with Oct4-bound sites which were determined as mentioned above. The co-shared regions were then annotated using annotatePeaks.pl script of HOMER. We have considered Intergenic co-shared regions only for OCT4 motif detection. Only sites with Oct4 motif were used for CRISPR/Cas9 targeting.

**RNA-Seq Analysis**

RNA-Seq libraries were mapped to mm9 genome using splice-aware STAR. Three or less mismatches were allowed in each read. The mapped libraries were then subjected to cuffdiff (Trapnell et al., 2012) to identify the differentially expressed genes and FPKM values of genes in different samples.

**Pearson correlation of RNA-seq**

The mapped RNA-Seq libraries were uploaded to SeqMonk v1.42.0 (https://www.bioinformatics.babraham.ac.uk/projects/seqmonk/). The pearson correlation was performed by using the PCA Analysis under the Data Store Similarity tab. The pearson values were then uploaded to R and heatmaps were generated using the heatmap.2 function of 'gplots'.

**PCA of RNA-seq**

The mapped RNA-Seq libraries were uploaded to SeqMonk v1.42.0 (https://www.bioinformatics.babraham.ac.uk/projects/seqmonk/). PCA was performed by using the Correlation Matrix under the Data Store Similarity tab.

**Unsupervised Clustering for differentially expressed gene**

The FPKM values of the significantly differentially expressed genes (vs NT) were used as a matrix. The matrix was uploaded to R and normalized per-row using the following formula (apply(data, 1, function(x)(x-min(x))/(max(x)-min(x)))).

The per-row normalized values were then used for generating the heatmap using heatmap.2 function in gplots.

**Lineage Enrichment Analysis CTEN analysis**

The list of differentially upregulated genes in each sample (vs NT) were uploaded to cTen (http://www.influenza-x.org/~jshoemaker/cten/) (Shoemaker et al., 2012) to identify the lineages in which these genes are significantly enriched.

**Reference**

Carpenter, A.E., Jones, T.R., Lamprecht, M.R., Clarke, C., Kang, I., Friman, O., Guertin, D.A., Chang, J., Lindquist, R.A., Moffat, J., et al. (2006). Genome Biol *7*, R100.

Chen, X., Xu, H., Yuan, P., Fang, F., Huss, M., Vega, V.B., Wong, E., Orlov, Y.L., Zhang, W., Jiang, J., et al. (2008). Integration of External Signaling Pathways with the Core Transcriptional Network in Embryonic Stem Cells. Cell *133*, 1106–1117.

Doench, J.G., Fusi, N., Sullender, M., Hegde, M., Vaimberg, E.W., Donovan, K.F., Smith, I., Tothova, Z., Wilen, C., Orchard, R., et al. (2016). Optimized sgRNA design to maximize activity and minimize off-target effects of CRISPR-Cas9. Nat. Biotechnol. *34*, 184–191.

Fang, H.T., EL Farran, C.A., Xing, Q.R., Zhang, L.-F., Li, H., Lim, B., and Loh, Y.-H. (2018). Global H3.3 dynamic deposition defines its bimodal role in cell fate transition. Nature Communications *9*, 1537.

Harms, D.W., Quadros, R.M., Seruggia, D., Ohtsuka, M., Takahashi, G., Montoliu, L., and Gurumurthy, C.B. (2001). Mouse Genome Editing Using the CRISPR/Cas System. Curr Protoc Hum Genet *83*:15.17.11-15.17.27.

Heinz, S., Benner, C., Spann, N., Bertolino, E., Lin, Y.C., Laslo, P., Cheng, J.X., Murre, C., Singh, H., and Glass, C.K. (2010). Simple combinations of lineage-determining transcription factors prime cis-regulatory elements required for macrophage and B cell identities. Molecular Cell *38*, 576–589.

Li, D., Liu, J., Yang, X., Zhou, C., Guo, J., Wu, C., Qin, Y., Guo, L., He, J., Yu, S., et al. (2017). Chromatin Accessibility Dynamics during iPSC Reprogramming. Cell Stem Cell *21*, 819–833.e6.

Mali, P., Moosburner, M., Church, G.M., Chari, R., and Church, G.M. (2015). Unraveling CRISPR-Cas9 genome engineering parameters via a library-on-library approach. *12*, 823–826.

Shoemaker, J.E., Lopes, T.J.S., Ghosh, S., Matsuoka, Y., Kawaoka, Y., and Kitano, H. (2012). CTen: a web-based platform for identifying enriched cell types from heterogeneous microarray data. BMC Genomics *13*, 460.

Stadhouders, R., Kolovos, P., Brouwer, R., Zuin, J., van den Heuvel, A., Kockx, C., Palstra, R.-J., Wendt, K.S., Grosveld, F., van IJcken, W., et al. (2013). Multiplexed chromosome conformation capture sequencing for rapid genome-scale high-resolution detection of long-range chromatin interactions. Nat Protoc *8*, 509–524.

Thongjuea, S., Stadhouders, R., Grosveld, F.G., Soler, E., and Lenhard, B. (2013). r3Cseq: an R/Bioconductor package for the discovery of long-range genomic interactions from chromosome conformation capture and next-generation sequencing data. Nucleic Acids Res. *41*, e132–e132.

Toh, C.-X.D., Chan, J.-W., Chong, Z.-S., Wang, H.F., Guo, H.C., Satapathy, S., Ma, D., Goh, G.Y.L., Khattar, E., Yang, L., et al. (2016). RNAi Reveals Phase-Specific Global Regulators of Human Somatic Cell Reprogramming. Cell Rep *15*, 2597–2607.

Trapnell, C., Roberts, A., Goff, L., Pertea, G., Kim, D., Kelley, D.R., Pimentel, H., Salzberg, S.L., Rinn, J.L., and Pachter, L. (2012). Differential gene and transcript expression analysis of RNA-seq experiments with TopHat and Cufflinks. Nat Protoc *7*, 562–578.

Zhang, Y., Liu, T., Meyer, C.A., Eeckhoute, J., Johnson, D.S., Bernstein, B.E., Nusbaum, C., Myers, R.M., Brown, M., Li, W., et al. (2008). Model-based analysis of ChIP-Seq (MACS). Genome Biol *9*, R137.
